# Quantifying Protein-Protein Interaction with a Spatial Attention Kinetic Graph Neural Network

**DOI:** 10.1101/2025.06.04.657832

**Authors:** Yuzhi Xu, Wei Xia, Chao Zhang, Xinxin Liu, Chengwei Ju, Xuhang Dai, Pujun Xie, Yuanqing Wang, Guangyong Chen, John Z.H. Zhang

## Abstract

Accurate prioritisation of near-native protein–protein interaction (PPI) models remains a major bottleneck in structural biology. Here we present SAKE-PP, a physics-inspired, spatial-attention equivariant graph neural network that directly regresses interface RMSD (iRMSD) without any native references. Trained on docking decoys generated through our novel hierarchical sampling strategy applied to PDB-Bind dataset, SAKE-PP combines force-field-like attention with Laplacian-eigenvector orientation to couple local interaction forces with global topology. On the 2024PDB benchmark comprising 176 heterodimers, SAKE-PP demonstrates effective optimization and selection of AF3 decoys, achieving improvements of 13.75% based on iRMSD statistics and 12.5% based on DockQ scores. It consistently outperforms the AF3 ranking score in multiple metrics, including overlap, hit rate, and correlation. In zero-shot evaluation of 139 antibody–antigen complexes, SAKE-PP improves the score–iRMSD correlation by 0.4. By unifying geometric deep learning with physics-based realism, SAKE-PP provides a robust, plug-and-play scoring function that streamlines reliable PPI evaluation and accelerates downstream structure-guided drug-design workflows.

## Introduction

Protein-protein interactions (PPIs) are fundamental to virtually all cellular processes, orchestrating complex biological functions from signal transduction and metabolic pathways to gene regulation. ^1–5^ Understanding the intricate structural details and energetic landscapes governing these interactions is crucial for deciphering the molecular mechanisms of life and for developing targeted therapeutic interventions

A paradigm shift in protein structure prediction has recently been catalyzed by deep learning methods, most notably AlphaFold and its subsequent iterations like AlphaFold-Multimer. ^7,8^ The latest version, AlphaFold3(AF3), has significantly expanded these capabilities, enabling accurate modeling of not only individual protein structures but also diverse biomolecular complexes, including protein-protein, proteinligand, and protein-nucleic acid assemblies. ^9^ To provide users with crucial information on prediction reliability, AlphaFold models output intrinsic confidence metrics such as predicted Local Distance Difference Test (pLDDT) for local accuracy, Predicted Aligned Error (PAE) for relative positioning, and interface Predicted Template Modeling (ipTM) score, specifically for assessing the quality of predicted interfaces in complexes. Training of these confidence metrics requires comparing predicted structures to known experimental data, an obvious question arises concerning the output of powerful prediction methods like AlphaFold: why haven’t they prioritized the direct prediction of these rigorous evaluation metrics, most notably iRMSD? Accurate estimation of iRMSD would be invaluable, as it would allow for the selection of predicted complex structures that are likely to be closer to the native state and thus serve as more suitable starting points for detailed downstream MD simulations. Deep-learning models including DeepBSP, RmsdXNA and DeepAccNet have demonstrated reference-free error regression for protein-ligand complexes or single-chain proteins, but the high-dimensional, interface-sensitive nature of PPI assemblies remains to be tackled. ^10–12^ The technical challenge of directly predicting such a metric likely stems from its inherent nature. Unlike the bounded nature of metrics such as pLDDT, PAE, and ipTM, iRMSD represents a precise, unbounded physical distance. ^13^ Accurately predicting such a continuous and potentially large value is significantly more challenging for current deep learning methods than predicting normalized confidence scores, posing a primary technical hurdle for directly modeling iRMSD.

In this paper, we propose a reference-free iRMSD predictor tailored to PPIs to bridge this gap by enabling Ångström-level scoring scoring of large decoy sets and seamless integration with physics-based refinement. This predictor, named SAKE-PP, is a novel physics-inspired spatial attention kinetics graph neural network(GNN) designed specifically for protein–protein evaluation. Unlike methods that rely on or are integrated within existing structure prediction architectures like AlphaFold’s PairFormer modules, SAKE-PP is designed as an independent model framework. This independent design is crucial to avoid any potential bias towards the specific outputs or feature representations of particular structure prediction models like AF3, ensuring the predicted iRMSD is a generalizable metric of geometric accuracy. SAKE-PP integrates two complementary modules to fully capture both structural and functional relationships at protein complex interfaces: the Spatial Attention Kinetics (SAKE) component simultaneously updates node features and their 3D coordinates, using a physics-inspired attention mechanism to account for spatial relationships among residues and simulate inter-residue forces under a force field; and the Laplacian Eigenvector Orientation component, which calculates the graph Laplacian eigenvectors to generate global embeddings reflecting each residue’s position within the overall fold, the steering message passing by the gradients and magnitudes of these eigenvectors to distinguish structurally similar but functionally distinct regions: finally, multiscale feature fusion unifies local interaction patterns with the global structural context to yield a holistic representation of the protein–protein interface. Training a model to accurately predict the precise, potentially unbounded values of iRMSD is a significant technical challenge. To construct a reliable training dataset for protein–protein docking evaluation, we curated 736 complexes from PDBBind and generated 1,472,000 candidate conformations using ZDock under a 15Ådistance restraint. A hierarchical sampling strategy was employed to mitigate the imbalanced iRMSD distribution of docking poses, with special emphasis on the critical 0–10 Årange. In particular, we selected representative and high-quality conformations through a combination of adaptive and uniform sampling. This procedure yielded a final dataset of 15,456 conformations, ensuring both diversity and enrichment of near-native structures to support robust regression-based model training.

Using this novel framework and training strategy, we demonstrate SAKE-PP’s superior performance and utility. Comprehensive benchmarking against state-of-the-art GNN-variants shows that our model achieves substantially lower RMSE and MSE, indicating superior accuracy in protein–protein interface prediction. Further, our model demonstrates effective optimization and selection of AF3 decoys on the newly curated 2024PDB dataset, achieving improvements of 13.75% based on iRMSD statistics and 12.5% based on DockQ scores. It consistently outperforms the AF3 ranking score in multiple metrics, including overlap, hit rate, and correlation. Moreover, as candidates progress along the drug-discovery pipeline, evaluation priorities shift towards developability attributes, notably energetic and dynamical properties such as binding affinity (Δ*G*_bind_) and kinetic stability (residence time and energetic barriers). ^14,15^ In this regime, our experiments favor the previous observation that RMSD-based geometric errors correlate well with total system energies obtained from MM/GBSA, free-energy perturbation (FEP) and related alchemical workflows (Fig. 1a). ^16,17^ A finely resolved iRMSD thus supplies a continuous, differentiable signal for selecting high-affinity, kinetically stable conformers and meshes naturally with downstream free-energy and molecular-dynamics analyses. Our model consistently avoids high-energy conformation selection errors frequently made by the AF3 ranking score and identifies a class of conformationally stable heterocomplexes. Furthermore, we performed antigen-antibody tests, which showed that despite no training on antigen–antibody complexes, SAKE-PP achieved a correlation score of 0.4 higher than that of the AF3 ranking score, indicating a strong generalizability. Ultimately, our development of an accurate, reference-free iRMSD predictor provides a crucial tool for evaluating and refining the structure of predicted protein complexes, thereby accelerating the mechanistic understanding and facilitating structure-guided design and discovery efforts in biology and medicine.

**Figure 1.**
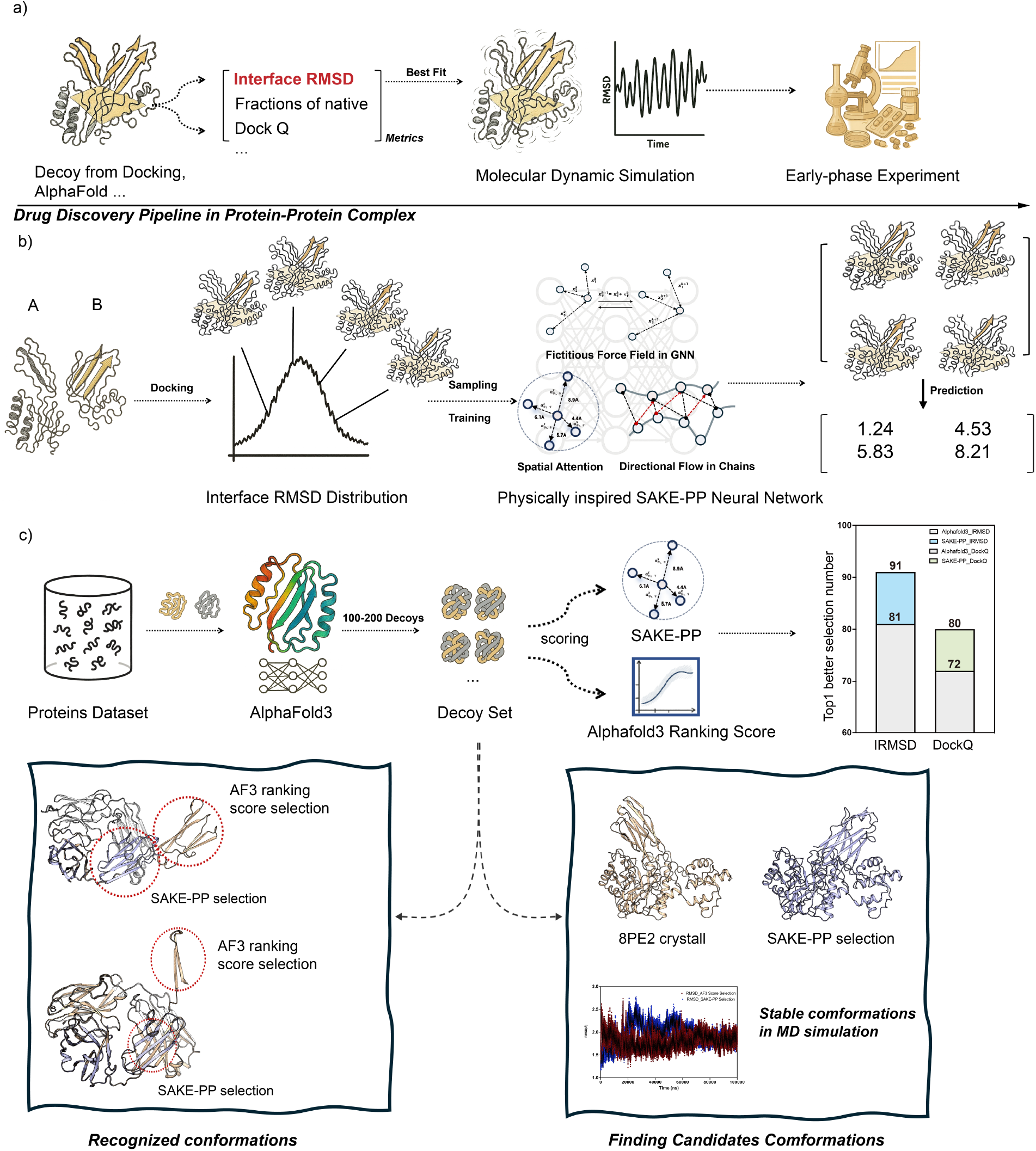
(a) In a typical protein–protein complex drug discovery pipeline, candidate structures are first generated via docking or AI-based prediction methods such as AF3. Each decoy is then evaluated using metrics such as interface RMSD(iRMSD), fraction of native contacts, or the composite DockQ score. Top-scoring conformations are subsequently subjected to all-atom molecular dynamics (MD) simulations and early-phase experimental validation. (b) SAKE-PP training workflow: Given a set of docking-generated complexes between proteins A and B, an iRMSD distribution is obtained, from which representative decoys are selected via hierarchical sampling strategies to train SAKE-PP. SAKE-PP is a physics-inspired, geometrically equivariant graph neural network that learns to predict iRMSD values for unseen decoys, guiding the identification of near-native binding modes. (c) Cross-dataset benchmarking and representative examples: For a diverse set of protein targets, AF3 is used to generate 100 decoys per target, which are then independently scored by SAKE-PP and AF3 ranking score. The bar chart shows the number of targets for which SAKE-PP (colored bars) or AF3 ranking score (gray bars) selects the top-1 decoy under iRMSD or DockQ. Bottom left: Comparison of SAKE-PP-selected low-energy conformations versus AF3 ranking score-selected high-energy, unstable alternatives for the same target. Bottom right: MD simulations of PDB 8PE2 over 1 *µ*s show that structurally distinct conformations selected by different scoring functions remain stable, suggesting the existence of multiple coexisting low-energy states in the protein–protein interaction landscape.

## Result

### Comparative Analysis of SAKE and AF3 Ranking Score in Protein-Protein Complex Structure Selection

AlphaFold 3 quantifies prediction reliability through a two-step process. First, it computes two complementary metrics: pLDDT (predicted Local Distance Difference Test, ranging from 0–100), which measures single-chain local folding accuracy, and ipTM (inter-chain predicted TM-score, ranging from 0–1), which evaluates the precision of multi-chain relative arrangements. These two metrics are then integrated into a unified scalar ranking score^1^, which enables the model to discriminate among a pool of structural decoys and identify the conformation it deems most reliable.

Although our model was initially trained using rigid-body decoys generated by ZDOCK, ^18^ we hypothesize that SAKE-PP effectively captured relevant structural features, allowing successful transfer to the more challenging task of flexible decoy discrimination, particularly in distinguishing AlphaFold-generated decoys.

For the protein docking tasks using AF3, we conducted a systematic docking analysis. The protein data primarily consisted of structures released in 2024, with a particular focus on those published after June 2024. Based on the FASTA sequence identifiers provided by the RCSB database, we performed a stringent selection of binary complex structures. ^19^ To be more specific, a selection protocol was implemented to curate binary complex structures: (1) exclusion of polypeptide chains containing fewer than 20 amino acid residues; (2) structural refinement using PDB4Fixer to remove crystallographic artifacts and resolve missing atoms; (3) systematic correction of noncanonical amino acids to their standard counterparts. As a result, we obtained a final dataset of 176 high-quality protein structures.

Based on the curated dataset of 176 high-quality protein structures, we constructed an end-to-end docking database using the AF3. Specifically, we used 20 random seeds of AF3 and generated 100 decoy conformations for each protein complex, thus creating a decoy database that captures a diverse conformational space. Furthermore, to evaluate the structural variability among these decoys, we computed and compared the iRMSD for the decoy structures of each protein complex. To enhance the robustness of the test set, we randomly selected 20% of the samples with receptor homologs as the test set, while the remaining 80% consisted of heterologous receptors. This partitioning strategy not only ensures that the test set covers samples with high receptor homology, but also evaluates the generalizability of the model in handling heterologous receptors.

### Overall Analysis

Figure 2 compares the effectiveness of selecting top-1 decoys based on the iRMSD and DockQ metrics between SAKE-PP and AF3’s ranking score. Among the 176 cases tested, SAKE-PP successfully identified better decoys based on iRMSD in 91 instances, whereas AF3 performed better in 81 instances, representing a 13.75% improvement for SAKE-PP. Notably, both methods selected identical top-1 decoys in four cases. Regarding DockQ, SAKE-PP achieved superior performance in selecting top-1 decoys in 80 instances compared to AF3’s 72 instances, indicating a 12.5% improvement. These results underscore the superior capability of SAKE-PP in effectively prioritizing high-quality structural decoys, highlighting its advantage over AF3 ranking score when evaluating AF3-generated protein complexes.

**Figure 2.**
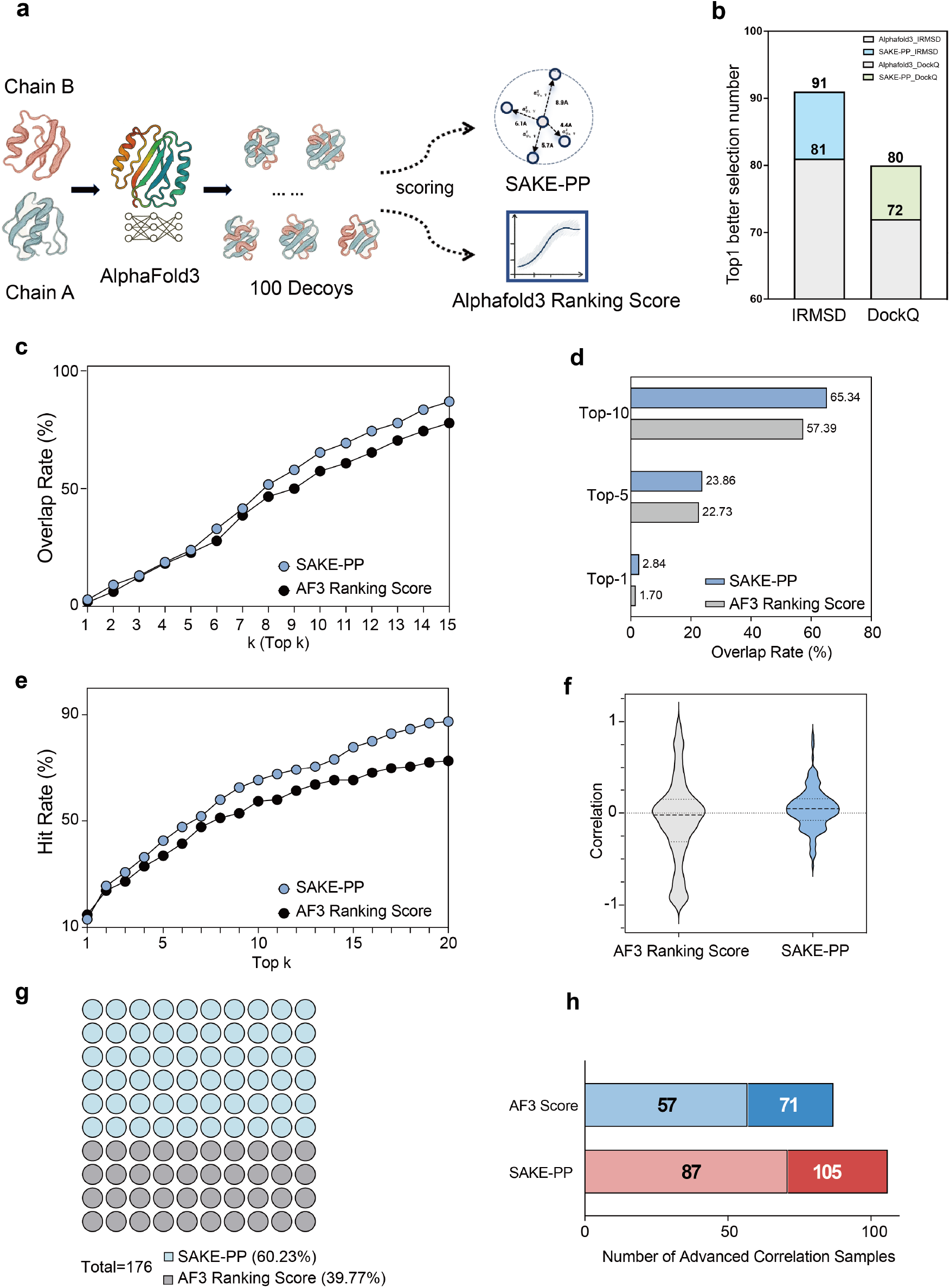
Benchmarking SAKE-PP against the AF3 ranking score on a 176-complex docking set. (a) Work-flow schematic. For each heterodimeric target, AF3 generates 100 decoys that are rescored independently by SAKE-PP and AF3 ranking score. (b) Top-1 selection quality. Bars indicate the number of complexes for which the Top-ranked decoy chosen by a given method exhibits the lower iRMSD (left) or the higher DockQ score (right). (c) Overlap-rate summarised at Top-1, Top-5, and Top-10. (e) Success-hit-rate curves: probability of finding at least one of the ten lowest-iRMSD decoys within the top-*k* ranks (1*≤ k ≤* 20). (f) Distribution of Pearson correlations (*r*) between predicted score and iRMSD for each complex. (g) Win–loss matrix (1 square = 1 complex). Blue squares denotecomplexes where SAKE-PP achieves a higher correlation; grey squares denote AF3 wins. SAKE-PP prevails in 105 of 176 cases (60.23%). (h) Win–loss bar: outer numbers show total wins (SAKE-PP 105, AF3 71); darker segment and inner numbers show wins with |Δ*r*| *>* 0.10 (SAKE-PP 87, AF3 57).

In addition to evaluating top-1 Beyond the top-1 assessment, we quantified the overlap between the top-*k* decoy lists produced by the two ranking schemes (Figure 2 c, d). Here, the *overlap rate* at a given *k* is simply the percentage of the 176 targets for which the top-*k* lists from SAKE-PP or AF3, when compared with the ground-truth ranking, share at least one common decoy. This metric gauges how consistently each method enriches structurally relevant candidates suitable for downstream molecular-dynamics (MD) refinement.

Across the entire top-k spectrum, SAKE-PP consistently outperforms AF3 ranking score. In the early stage (*k* = 1–3), it delivers a 67% relative gain at Top-1 and maintains a clear edge up to *k* = 5, ensuring high precision among the highest-ranked decoys. From *k* = 6 onward the advantage widens, with absolute gains exceeding 5 percentage points and relative improvements peaking at 18% (*k* = 6) and remaining *≥* 14% through *k* = 12, evidencing SAKE-PP’s ability to recover high-quality decoys overlooked by AF3 ranking score. At the upper bound (*k* = 15) SAKE-PP achieves an 86.93% overlap versus 77.84% for AF3 ranking score—a 9.09 pp improvement—and the two curves never intersect, indicating globally superior ordering. Collectively, these results demonstrate SAKE-PP’s strong early enrichment and sustained superiority across broader decoy pools, making it highly suitable for ensemble-based free-energy or MD workflows that require multiple accurate starting structures.

Besides, following the CAPRI convention, we designate the ten decoys with the lowest iRMSD for each complex as reference “near-native” poses—an operational choice that circumvents the difficulty of unambiguously identifying true native structures within a decoy set. ^20^ Accordingly, the success-hit-rate is the probability of retrieving at least one of these references within the top-*k* ranks. As illustrated in Figure 2e, SAKE-PP outperforms the AF3 ranking score from *k* = 2 onward. Averaged over *k* = 1–5, the success-hit rate is 29.7% versus 27.2% (+9.2% relative). In the *k* = 6–20 range the gap widens to 71.1% versus 61.2% (+16.2%). Across the full interval (*k* = 1–20), SAKE-PP attains a mean success-hit rate of 60.7%, exceeding AF3 ranking score by 8.0 percentage points (+15.3%). The practical screening depth of *k* = 20 benefits most, showing a 14.8 pp lift (87.5% vs 72.7%, +20%). Statistical analysis confirms the robustness of this improvement (paired *t* = 7.83, *p ≈* 2.3 *×* 10^*−*7^; Cohen’s *d* = 1.75, large effect). These results highlight the superior discriminative power of SAKE-PP in prioritising near-native decoys—especially when workflows allow more than a single decoy to be forwarded for downstream refinement.

To complement the early-ranking evaluation, we examined how well each scoring function tracks overall decoy quality. For every target we computed the Pearson correlation coefficient (*r*) between its raw score and iRMSD. Head-to-head tallies (Figure 2g) indicate that SAKE-PP yields a higher *r* in 105 of 176 cases (60.23%), whereas the AF3 ranking score prevails in 71 cases (39.77%), corresponding to a 51.4% relative improvement in win–loss ratio. To focus on meaningful differences, we further required |Δ*r*| *>* 0.10 (“advanced” threshold). Under this stricter criterion (Figure 2h) SAKE-PP still surpasses AF3 ranking score in 87 complexes, while lagging in 57, reaffirming the robustness of the advantage. A two-sided Wilcoxon signed-rank test confirms statistical significance (*Z* = 7.35, *p ≈* 1.9 *×* 10^*−*13^).

### Successful Cases

To further demonstrate the practical utility of SAKE-PP, we turned to three challenging yet representative complexes—8S4K, ^21^ 8SOZ, ^22^ and 8VGG, ^23^ the conformations directly selected by AF3 ranking score yielded iRMSD values of 17.886Å, 16.352Å, and 4.719Å, respectively, all significantly exceeding the acceptable threshold of 4.0 Å. Notably, this discrepancy does not stem from structural inaccuracies of AF3-generated decoys themselves; rather, SAKE-PP successfully identified near-native conformations from these AF3 decoys, achieving markedly improved iRMSD values of 1.521Å, 1.885Å, and 0.925Å, respectively. This highlights the effectiveness of SAKE-PP in accurately distinguishing conformations closely resembling native structures.

Specifically, among the decoys generated by AF3, the iRMSD values exhibit a distinct bimodal distribution. Approximately 87% of the predictions cluster around a high average iRMSD of 17.26 Å (designated as the “deviated cases” group), while only about 13% cluster around a low average iRMSD of 1.526 Å (defined as the “reasonable cases” group). The overall mean iRMSD across all predictions is 15.22 Å.

As shown in Figure 2(d), the decoy selected by AF3 ranking score belongs to the deviated group, whereas SAKE successfully identifies a decoy from the reasonable group. A notable positional displacement at a key structural region differentiates these two conformations. Further analysis of the AF3 ranking scores reveals that the decoy selected by AF3 ranking score possesses the highest ranking score (0.8709), while the SAKE-selected decoy receives a lower AF3 ranking score of 0.8385.

To investigate whether AF3 ranking score’s selection of the deviated conformation was incidental or indicative of a general preference, we randomly sampled ten decoys from each group for a detailed analysis. For the deviated group, we found that the high iRMSD values primarily stem from substantial displacements of predicted structures relative to the binding site. AF3 ranking score consistently assigned these displaced decoys high ranking scores, averaging 0.8572 (standard deviation: 0.005). Conversely, the randomly sampled decoys from the reasonable group exhibited significantly lower AF3 ranking scores, averaging 0.8399 (standard deviation: 0.004). This suggests that the AF3 ranking score function intrinsically favors the deviated cases.

Interestingly, we also observed one exceptional case within the reasonable group: a structure with a low iRMSD of 1.495 Å obtained a relatively high AF3 ranking score of 0.8501. Although this score remains below the majority of deviated cases, it is still higher than the average score of the reasonable group. This result indicates that the AF3 ranking score does not effectively differentiate subtle but critical structural deviations at the binding site.

In contrast, the top seven conformations ranked by SAKE-PP all fall within the reasonable structural range, indicating the robustness of its predictive performance.

Similar to the case of 8S4K, most decoys for 8VGG fall within the near-native range, with *≈* 77% exhibiting iRMSD *<* 2 Å. Nevertheless, about 23% of the structures show iRMSD values at or above 4 Å. These high-error outliers arise from the same issue observed in 8S4K—namely, the presence of a solvent-exposed extended loop/*β*-strand whose position is difficult to model accurately. Interestingly, even though 77% of the ensemble represents near-native conformations, the AF3 ranking score still assigns disproportionately high weight to the erroneous decoys.

The case of 8SOZ is more extreme: the best conformation ranks only 7^th^ out of 100 based on the AF3 ranking score, despite having a near-native RMSD of 1.885 Å and representing just 1% of the ensemble. In contrast, the remaining 99% of conformations show substantial deviations, with an average RMSD of 16.5 Å. Structural comparison suggests that this discrepancy is similarly caused by an anomalous domain-level deviation.

Notably, the RMSDs of the six decoys ranked in AF3 ranking score before and after the best one are all around 16.5 Å. As shown clearly in Figure 3e, SAKE-PP is able to accurately identify this extreme near-native structure from the 100 conformations. This demonstrates SAKE’s potential as a more effective scoring function that can complement the limitations of AF3. In cases with large structural heterogeneity, the AF3 rank score fails to discriminate between high- and low-quality models, and the scores it assigns do not reflect the actual differences in structural quality. In contrast, SAKE-PP assigns a score of 1.268Åto the best conformation, while its predictions for all other conformations with higher iRMSD values are consistently above 4Å.

**Figure 3.**
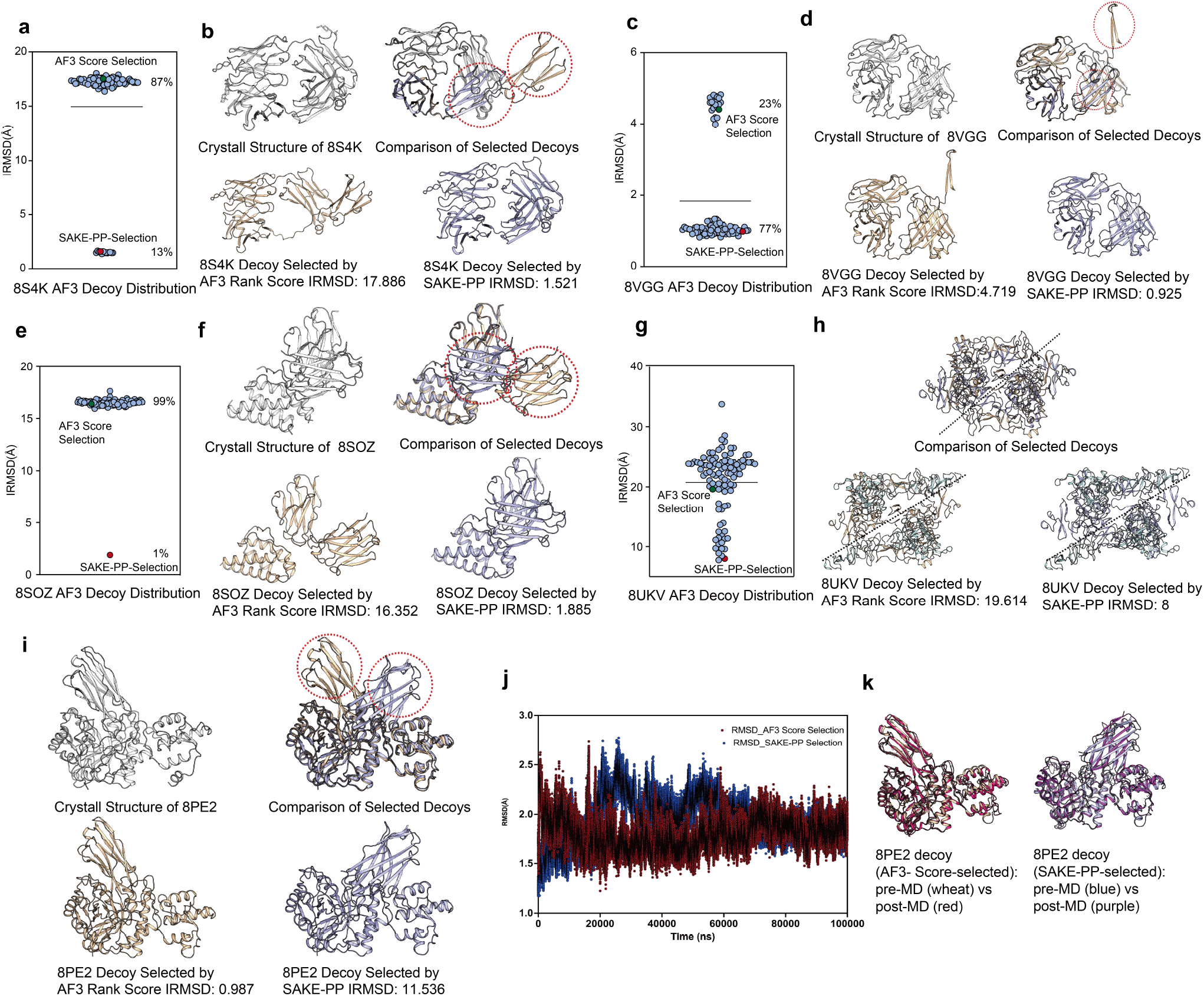
(a, c, e, g) Scatterplots of iRMSD for all 100 AF3-generated decoys of PDB IDs 8S4K, 8VGG, 8SOZ, and 8UKV. The symbol highlighted in red marks the decoy chosen by SAKE-PP; the blue symbol marks that chosen by the AF3 ranking score. Percentages indicate the fraction of decoys with equal or lower iRMSD than the AF3 ranking score-selected model. (b, d, f, h) Structural overlays. For each target the crystal structure is shown in grey. Decoys selected by AF3 ranking score (tan) and SAKE-PP (slate-blue) are superposed; red dashed circles highlight regions where the two selections differ markedly. Corresponding iRMSD values are listed beneath each panel. (i) Same overlay for 8PE2, where SAKE-PP intentionally chose a more dissimilar starting pose (iRMSD = 11.536 Å) to test sampling robustness. (j) All-atom MD refinement of the AF3 ranking score-selected (wheat) and SAKE-PP-selected (blue) 8PE2 decoys: backbone RMSD vs simulation time over 1 *µ*s. Post-MD conformations are coloured red (AF3) and purple (SAKE-PP). (k) Final MD snapshots of the two 8PE2 decoys, showing that the SAKE-PP start converges towards the experimental structure while the AF3 ranking score start drifts away.

As shown in Figures 2e and f, this is similarly a case where scoring bias arises from an outlier conformation with abnormal structural deviations. Although the proportion of such deviated structures is 23%, AF3 ranking score still selected one of the deviated conformations as the top model. In contrast, SAKE-PP once again demonstrated robustness in handling such cases by correctly identifying the near-native structure.

### Failure Cases

Although SAKE-PP generally performs well, we identified several high-ranking cases where its predictions are still suboptimal. Closer inspection shows that these errors all arise from pose-selection issues.

In the 8PE2 ^24^ complex evaluation, the decoy selected by SAKE-PP (iRMSD = 11.536 Å) exhibits global homology to the structure chosen by AF3 ranking score (iRMSD = 0.987 Å). Despite a substantial C_*α*_-RMSD difference, both models superimpose remarkably well in the central flexible loop and C-terminal *β*-sheet regions. Detailed structural comparison reveals that the *β*-sheet domain (residues 444–561) adopts a mirror-like orientation: the normals of their sheet planes differ by approximately 130^*°*^, indicating pronounced reorientation while preserving the core fold architecture. Notably, the intra-sheet hydrogen-bond network and local backbone geometry remain intact, emphasizing the conformational stability of this region.

Each decoys was solvated in an explicit TIP3P water box containing 0.15 M NaCl and subjected to 1 *µ*s all-atom MD simulations using the Amber ff14SB force field at 300 K and 1 atm. The potential energy converged within the first 200 ns and stabilized at approximately *−*8.0 *×* 10^5^ kJ mol^*−*1^, with subsequent fluctuations of no more than ±5 *×* 10^3^ kJ mol^*−*1^. Throughout the simulation trajectory, secondary structure content varied by less than ±2%, the hydrogen-bond pattern remained unchanged, and the solvent-accessible surface area (SASA) deviated by under 8%. B-factor analysis further demonstrates that the highly mobile regions in the SAKE-PP decoy coincide with those in the AF3 ranking score-selected structure, highlighting their shared dynamic flexibility profile.

To further dissect the sources of error, we examined three representative complexes—8XYZ, 8YF2, and 8K0E (see Fig.S1)—in which SAKE-PP performed notably worse while AF3 ranking score yielded highly accurate predictions. In all three cases, the conformations selected by SAKE-PP deviated from the crystal structures by more than 16 Åin terms of iRMSD, whereas the AF3-selected models consistently stayed within 2 Å. Interestingly, both predictions showed nearly identical backbone folds, suggesting that the discrepancy arose primarily from differences in the relative positioning of the ligand or partner chain within the binding pocket.

In particular, for complexes 8XYZ and 8YF2, SAKE-PP’s selection tends to push the ligand deeply into a hydrophobic pocket formed by a contracted flexible loop, aiming for tight van der Waals packing and geometric complementarity. On the other hand, AF3 ranking score selection decoy adheres more closely to biologically informed configurations observed in signaling proteins—even if these poses are not always the most sterically complementary. From a structural standpoint, AF3 ranking score’s predictions may sacrifice some geometric fit in favor of recapitulating the native binding mode more faithfully. For 8K0E, both SAKE-PP and AF3 insert the ligand into a deep pocket formed by two distinct regions of the protein. However, SAKE-PP’s chosen pocket is defined by two *β*-strands that pack very closely, promoting a more compact interface, whereas AF3 selects a pocket that facilitates broader, larger-scale contacts.

While our previous analysis primarily relied on geometric configurations and statistical model outcomes to distinguish successful from failed cases—an approach that inherently carries certain biological information constraints—we were equally interested in examining these cases from a physical energetics standpoint. Accordingly, we performed molecular dynamics simulations and computed the corresponding binding energies. Remarkably, the binding energy calculations consistently revealed that decoys selected by SAKE-PP demonstrate energetic superiority over those identified by AF3 ranking score, displaying enhanced binding affinities.

Across all three systems, the stable conformations obtained by SAKE-PP exhibited higher proteinprotein binding energies than those selected by AF3 ranking score. As Figure S2, specifically, in the 8K0E-AF3 ranking score system, there were 144 residues from chain A within a 10 Å range of chain B, more than the 116 residues in 8K0E-SAKE-PP, with 113 residues appearing in both models. Although AF3 ranking score obtained an additional 31 residues contributing a total of -1.78 kcal/mol, SAKE-PP’s predicted structure showed a binding energy 4.36 kcal/mol stronger than AF3 ranking score, indicating that within the same residues predicted by both models, the dynamic simulation conformations starting from SAKE-PP’s predicted structure exhibited stronger binding to chain B. As Table S1 shown, residues such as 92ARG, 236GLN, 96PRO, 65PRO, 83LEU, 61TRP, 231ASN, 55TRP, 229GLU, 69SER, 235PHE, and 46ARG contributed more than 1 kcal/mol stronger in SAKE-PP compared to AF3 ranking score. These residues were involved in interactions mainly related to pi-pi interactions, pi-cation interactions, pi-alkane interactions, and hydrogen bonding. On the other hand, residues 467GLU, 644ASP, 632ASP, and 636MET obtained only in the AF3 ranking score model provided repulsive sites with energies exceeding 1 kcal/mol, primarily due to electrostatic repulsion.

Both models identified 35 hotspot residues. Therefore, the conformations obtained by 8K0E-SAKE-PP strengthened the interactions between hotspot residues at the interface. Despite the addition of multiple binding sites in 8K0E-AF3 ranking score, the presence of evident repulsive sites in these locations not only failed to introduce new significantly enhanced binding regions but also weakened binding in other areas.

In the 8YF2-SAKE-PP system, there were 86 interface residues, compared to 59 in 8YF2-AF3 ranking score, resulting in an additional 27 residues (Figure S3) contributing -9.21 kcal/mol to the binding energy. Notably, residue 17TYR contributed 4.44 kcal/mol, while 12ARG contributed 3.38 kcal/mol. Positively charged residues such as 175ARG, 90LYS, 132ARG, and 21ARG all contributed over 2 kcal/mol, while the polar residue 136THR contributed 0.99 kcal/mol. These additional residues obtained by SAKE-PP provided new binding sites and regions for binding.

Furthermore, among the interface residues common to both SAKE-PP and AF3 ranking score, residue 83LYS exhibited the largest increase, shifting from -2.36 kcal/mol in AF3 ranking score to -7.05 kcal/mol in SAKE-PP. Residue 87TYR changed from a repulsive contribution of 0.21 kcal/mol in AF3 ranking score to a positive contribution to binding of -3.28 kcal/mol in SAKE-PP, indicating the contribution of the pi-system interactions. Additionally, residues 138ILE, 156PHE, and 159GLN showed increases in contribution exceeding 2 kcal/mol. Overall, the combined contributions of these residues in SAKE-PP not only introduced new binding sites but also enhanced the effects of overlapping residues (15.38 kcal/mol), with electrostatic interactions and pi-system interactions playing significant roles.

These results indicate that SAKE-PP exhibits a strong correlation with binding energy, consistently favoring decoy conformations with more favorable binding affinities.

### Antibody:SAKE-PP benchmark beyond the training distribution

To probe how well SAKE-PP performs outside the distribution on which it was trained, we designed an antibody–antigen benchmark. This setting is deliberately challenging: the model was optimized only on a generic set of dimeric protein–protein interactions and has never seen multi-chain antibody–antigen complexes during training. Its performance here therefore provides a direct measure of its ability to generalize to entirely new protein classes.

For the benchmark we began with the SAbDab dataset curated by Fang et al. Redundancy was removed, chains shorter than 20 residues were discarded, and only entries containing complete antibody–antigen pairs were retained. ^25,26^ After these filters, 139 unique antibody–antigen complexes remained and were used for evaluation. For each complex, AF3 first generated 200 decoys, dnd then this decoy set was re-ranked by SAKE-PP(Fig. 4a). Globally, the Pearson correlation between score and iRMSD is *−*0.2886 for AF3 ranking score indicating a pronounced negative trend—whereas SAKE-PP improves the correlation to +0.1116, a gain of almost +0.40. On a per-complex basis, SAKE-PP outperforms AF3 ranking score in 106 of 139 cases (76.3%), while AF3 ranking score is superior in only 33 cases (23.7%). Strikingly, 70 complexes (*>*50% of the test set) flip from negative to positive correlation when rescored with SAKE-PP, underscoring the method’s stronger interface-aware ranking capability.Although SAKE-PP trails the native AF3 ranker by a single complex in the “pick-one” headline metric (AF3 ranking score selects the lower-iRMSD top-1 decoy in 70 cases, SAKE-PP in 69), a closer look at the difficult subset reveals a very different picture. We isolated the 18 antibody–antigen complexes for which AF3 ranking score assigns a high confidence score (ranking score > 0.75) yet the corresponding decoy is clearly wrong (iRMSD > 4 Å). In other words, these are the instances where the AF3 score is most misleading. Within this challenging cohort, SAKE-PP outperforms AF3 ranking score in 14 out of 18 cases, correctly demoting the erroneous high-score decoys and surfacing nearer-native alternatives.

**Figure 4.**
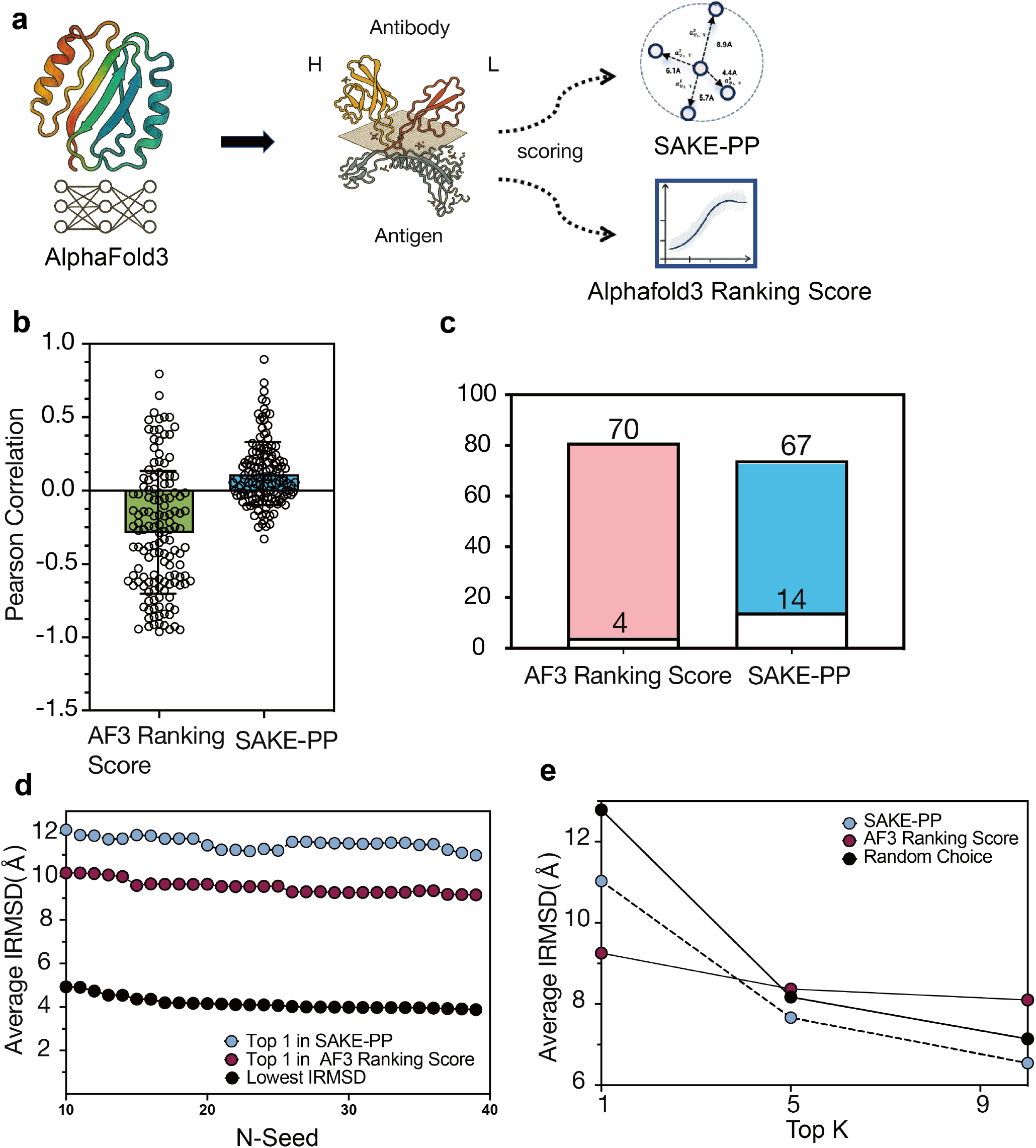
Antigen benchmark: SAKE-PP versus the AF3 ranking score. (a) Workflow. AF3 first generates 100 decoys for each heavy-/light-chain antibody bound to its antigen; the decoys are then rescored by SAKE-PP (blue) or by the native AF3 ranking score (grey). (b) Pearson correlations (*r*) between predicted score and iRMSD for every complex (*n* = 74). Each point is a single complex; boxes indicate the inter-quartile range. AF3 correlations cluster around negative values (green), whereas SAKE-PP correlations are predominantly positive (blue). (c) Count of complexes with positive versus negative correlation. AF3 ranking score yields only 4 positive and 70 negative cases, while SAKE-PP reverses the trend with 67 positive and 14 negative cases. (d) Seed-selection experiment. As the number of seeds (*N*) increases, the average iRMSD of the single best seed chosen by SAKE-PP (blue) or by AF3 ranking score (purple) is compared with the theoretical optimum—the lowest-iRMSD decoy in the pool (black). (e) Average iRMSD achieved when retaining the top-*k* decoys (*k* = 1, 5, 9). SAKE-PP (blue) consistently outperforms AF3 ranking score (purple) and a random baseline (black dashed). 30

Abramson et al. have argued that attaining peak accuracy with AF3 in antibody-antigen system requires generating and re-ranking a very large number of seed decoys, which incurs substantial computational overhead. ^9^ To test this claim, we expanded each antibody–antigen decoy pool from 50 to 200 structures and retained only the top-ranked model according to either SAKE-PP or the native AF3 ranking score, while also recording the theoretical optimum (the lowest iRMSD in the pool). As shown in Fig. 4e, the mean iRMSD of all three traces decreases as the number of seeds doubles, confirming that deeper sampling indeed increases the chance of encountering near-native poses. AF3 ranking score’s average remains slightly better than that of SAKE-PP, a difference driven mainly by a few outlier complexes (e.g., 7XJ6, 7XJ8) where SAKE-PP makes large orientation errors—highlighting that the method can still be misled by extreme local heterogeneity.

From a downstream perspective, however, what matters most is the quality of the top-*k* candidates that will proceed to molecular-dynamics refinement or affinity maturation. We therefore compared the best iRMSD found within the top 1, top 5, and top 10 decoys out of 200 (Fig. 4f) and benchmarked them against randomly chosen sets of the same size. Across all thresholds, SAKE-PP consistently outperforms the random baseline and converges to an average iRMSD of *∼*6 Å for the top 10, whereas AF3 ranking score lags behind random choice in the top-5 and top-10 scenarios, indicating that its native score struggles to enrich high-quality structures in large decoy pools. Taken together, while more extensive sampling benefits both methods in principle, SAKE-PP is markedly better at concentrating near-native decoys toward the very top of the ranking, delivering a smaller yet higher-quality candidate set that keeps subsequent MD simulations both reliable and computationally manageable.

It is important to emphasize that SAKE-PP was never trained on antibody–antigen complexes; it was optimized solely on a general protein–protein dataset. The strong performance observed here therefore underscores the model’s ability to generalize beyond its training distribution.

### Ablation Study

We conducted an ablation study to evaluate the impact of our hierarchical iRMSD-guided sampling strategy. SAKE-PP employs uniform sampling across the full interface-RMSD distribution, while SAKE-PP-ns follows conventional protocols by selecting only the top 100 decoys per complex ranked by docking score. Five-fold cross-validation revealed substantial improvements with hierarchical sampling: mean absolute error decreased from 3.45 ± 0.01 to 0.95 ± 0.02 (73% reduction), root-mean-square error dropped from 4.34 ± 0.01 to 1.29 ± 0.03 (70% reduction), and Pearson correlation coefficient increased from 0.24 ± 0.03 to 0.62 ± 0.01 (156% improvement). These results demonstrate that hierarchical sampling enables more robust structure-property relationships and superior ranking consistency.

Training dynamics further confirm these performance gains(Fig.S4 and Fig.S5). SAKE-PP-ns exhibited slow convergence, with training loss decreasing gradually from 26 to 18 over 20 epochs while validation loss fluctuated between 18-20, suggesting potential overfitting. Conversely, SAKE-PP with hierarchical sampling achieved rapid convergence, dropping from 4.5 to 1.6 within 10 epochs, with validation loss closely tracking training loss and minimal cross-validation variance. The synchronized decline in both training and validation metrics, coupled with effective learning rate scheduling, indicates superior generalization and reduced sensitivity to data partitioning. These training characteristics corroborate the quantitative improvements and demonstrate enhanced optimization stability conferred by hierarchical iRMSD-guided sampling.

The comparative performance of SAKE-PP and baseline models under 5-fold and 10-fold cross-validation is summarized in Table 3 and visually depicted in Figure 5. Our proposed model demonstrates consistently superior performance across all evaluation metrics. In the 5-fold cross-validation setting, SAKE-PP achieves a mean MAE of 0.9451 (±0.0157), significantly outperforming EGNN ^27^ (0.9922 ±0.0203), DGN ^28^ (0.9966 ±0.0796), and traditional GNN architectures such as GCN ^29^ and GAT^30^ (Figure 5a). Similar performance advantages are observed for RMSE (1.2912 ±0.0275 vs. 1.2986–1.9126, Figure 5b) and Pearson’s R (0.6164 ±0.0118 vs. *≤*0.6035, Figure 5e), confirming both higher predictive accuracy and stronger correlation with ground truth values. Notably, SAKE-PP exhibits greater stability, evidenced by consistently lower standard deviations across all evaluated metrics. For instance, under 5-fold validation, SAKE-PP’s MAE standard deviation (0.0157) is approximately 22% smaller compared to EGNN (0.0203) and remarkably 80% smaller relative to DGN (0.0796). Such robustness is equally apparent in the 10-fold validation scenario, where SAKE-PP maintains a narrower MAE spread of 0.0336 versus DGN’s 0.0932 (Figure 5c), indicating superior generalization and less sensitivity to data splits. The RMSE metric further supports these findings, with SAKE-PP showing reduced variance and consistently lower mean errors (Figure 5d). Overall, these analyses underscore SAKE-PP’s superior predictive performance and generalization capabilities compared to existing baseline models, highlighting its potential applicability to broader and more varied datasets.

**Table 1:** Comparison of Predicted Binding Free Energies Using AF3 Ranking Score and SAKE-PP Methods.

| PDB ID | Method | Binding Free Energy (kcal/mol)↓ |
| --- | --- | --- |
| 8K0E | AF3 ranking score | -71.41 |
| 8K0E | SAKE-PP | **-75.77** |
| 8YF2 | AF3 ranking score | -26.39 |
| 8YF2 | SAKE-PP | **-50.99** |
| 8XYZ | AF3 ranking score | -29.39 |
| 8XYZ | SAKE-PP | **-30.97** |

**Table 2:** Impact of Hierarchical iRMSD-Guided Sampling on Model Performance (mean ±} Std)

| Model | MAE | RMSE | R (Corr.) |
| --- | --- | --- | --- |
| Baseline (No Sampling) | 3.4477 ± 0.0089 | 4.3445 ± 0.0145 | 0.2411 ± 0.0342 |
| **Our Model** (Hierarchical Sampling) | **0.9451** ± 0.0157 | **1.2912** ± 0.0275 | **0.6164** ± 0.0118 |
| **Improvement (%)** | 72.59% ↓ | 70.28% ↓ | +155.66% ↑ |

**Table 3:** Model Comparison under 5-Fold and 10-Fold Cross-Validation. The results are presented as mean ± standard deviation (Std). **Bold** indicates the best performance and underline indicates the second-best.

| 5-Fold | MAE | RMSE | R |
| --- | --- | --- | --- |
| Our Model | **0.9451** ± 0.0157 | <u>1.2912</u> ± 0.0275 | **0.6164** ± 0.0118 |
| EGNN<sup>27</sup> | <u>0.9922</u> ± 0.0203 | 1.2986 ± 0.0555 | 0.5983 ± 0.0071 |
| DGN<sup>28</sup> | 0.9966 ± 0.0796 | **1.2886** ± 0.2507 | <u>0.6035</u> ± 0.0810 |
| GCN<sup>29</sup> | 1.0983 ± 0.0186 | 1.423 ± 0.0490 | 0.4933 ± 0.0129 |
| GAT<sup>30</sup> | 2.0448 ± 0.2599 | 1.9126 ± 0.3420 | 0.0021 ± 0.0177 |
| HGT<sup>31</sup> | 1.2081 ± 0.0179 | 1.575 ± 0.0066 | 0.0059 ± 0.0240 |
| 10-Fold | MAE | RMSE | R |
| Our Model | **0.9466** ± 0.0336 | **1.2639** ± 0.0357 | **0.6240** ± 0.0206 |
| EGNN | <u>0.9892</u> ± 0.0292 | <u>1.2705</u> ± 0.1119 | 0.6034 ± 0.0244 |
| DGN | 0.9860 ± 0.0932 | 1.2787 ± 0.3093 | <u>0.6094</u> ± 0.0831 |

**Figure 5.**
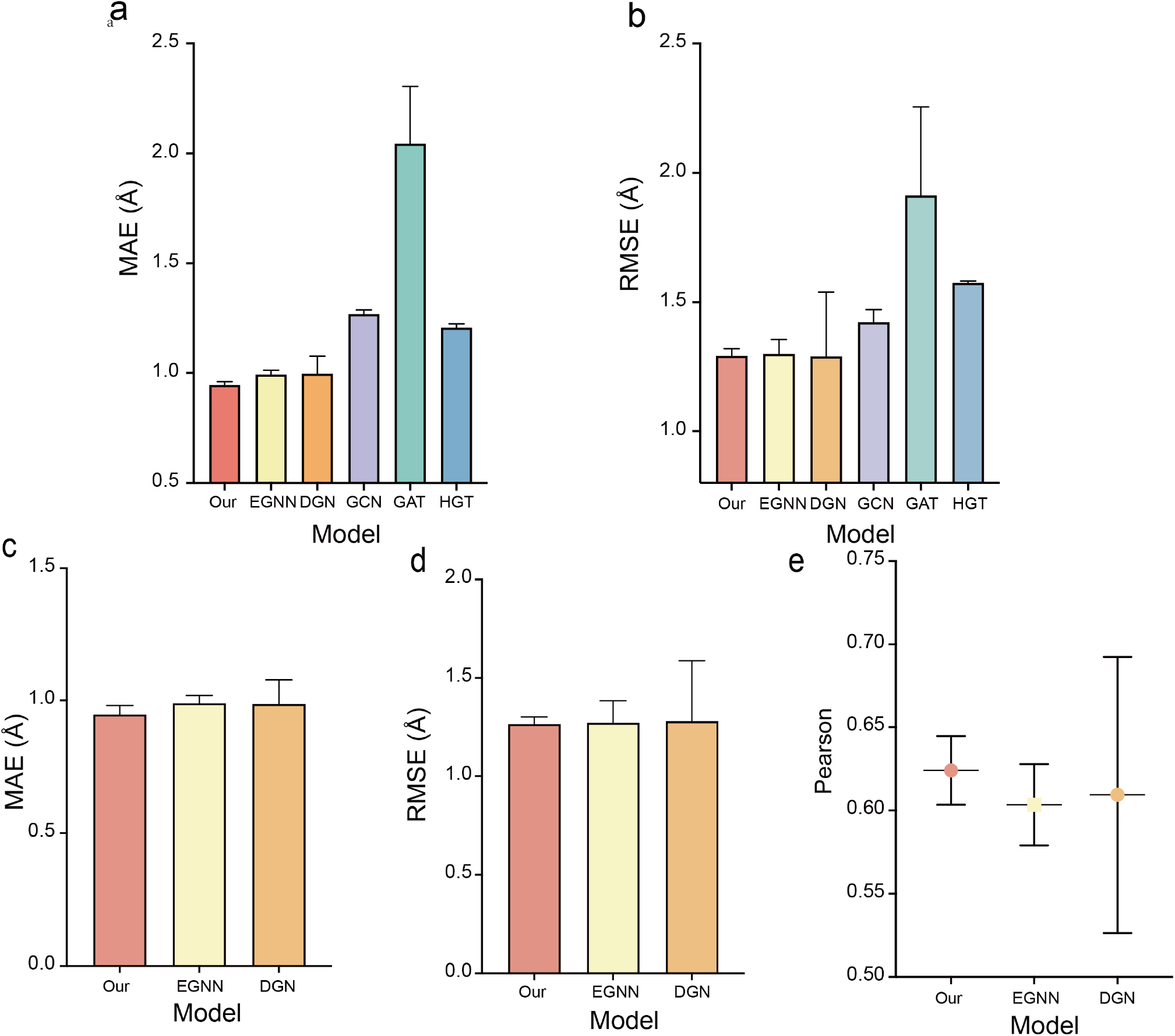
Comparative performance analysis of SAKE-PP and baseline GNN models under 5-fold (a, b) and 10-fold (c–e) cross-validation. Panels (a) and (b) depict Mean Absolute Error (MAE) and Root Mean Square Error (RMSE), respectively, comparing SAKE-PP against EGNN, DGN, GCN, GAT, and HGT in the 5-fold setting. Panels (c–e) illustrate the performance metrics (MAE, RMSE, and Pearson correlation coefficient, respectively) under 10-fold cross-validation. (b) Conformational distribution after sampling. By CAPRI criteria (iRMSD*≤* 4 Å vs. *>* 4 Å), 42.87% of decoys are acceptable (light blue) and 57.13% are unacceptable (peach); within the acceptable set, high-quality poses (0–2 Å; 34.01%, red) and borderline poses (2–4 Å; 65.99%, blue) are shown. (c) iRMSD distribution of all 1,472,000 decoys in the 0–10 Å range, showing a “sandglass” shape with enrichment at high RMSD. (d) iRMSD distribution after sampling, illustrating balanced coverage and enrichment in the critical 0–4 Å region.

**Figure 6.**
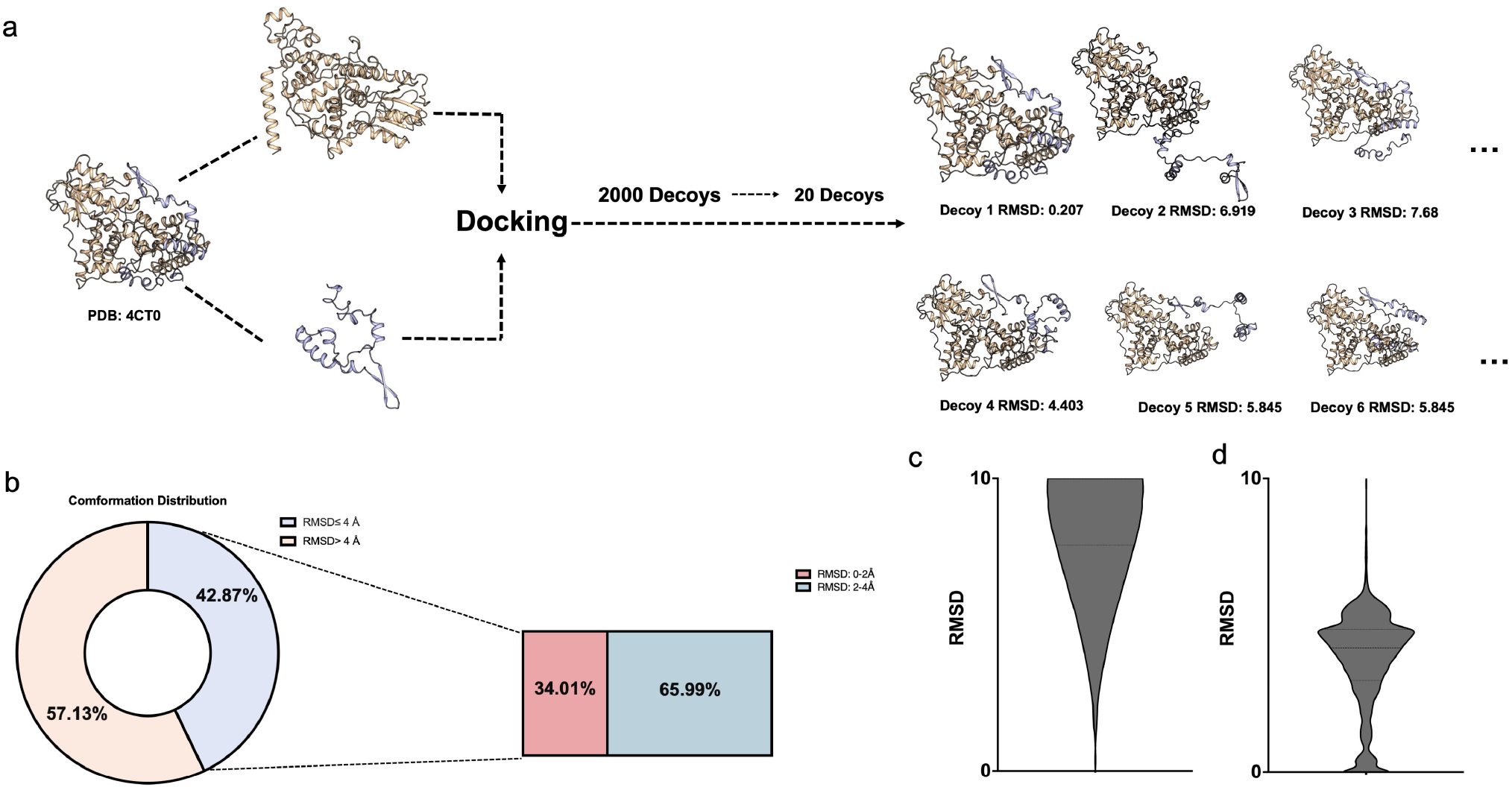
**(a)** From PDBBind, 736 protein–protein complexes were docked with ZDock under a 15 Å distance restraint to generate 2,000 decoys each, then hierarchically sampled down to 20 representative poses. Right: representative decoys for PDB 4CT0 and their iRMSD values. **(b)** Conformational distribution after sampling. By CAPRI criteria (iRMSD *≤* 4 Å vs. *>* 4 Å), 42.87% of decoys are acceptable (light blue) and 57.13% are unacceptable (peach); within the acceptable set, high-quality poses (0–2 Å; 34.01%, red) and borderline poses (2–4 Å; 65.99%, blue) are shown. **(c)** iRMSD distribution of all 1,472,000 decoys in the 0–10 Å range, showing a “sandglass” shape with enrichment at high RMSD. **(d)** iRMSD distribution after sampling, illustrating balanced coverage and enrichment in the critical 0–4 Å region.

**Figure 7.**
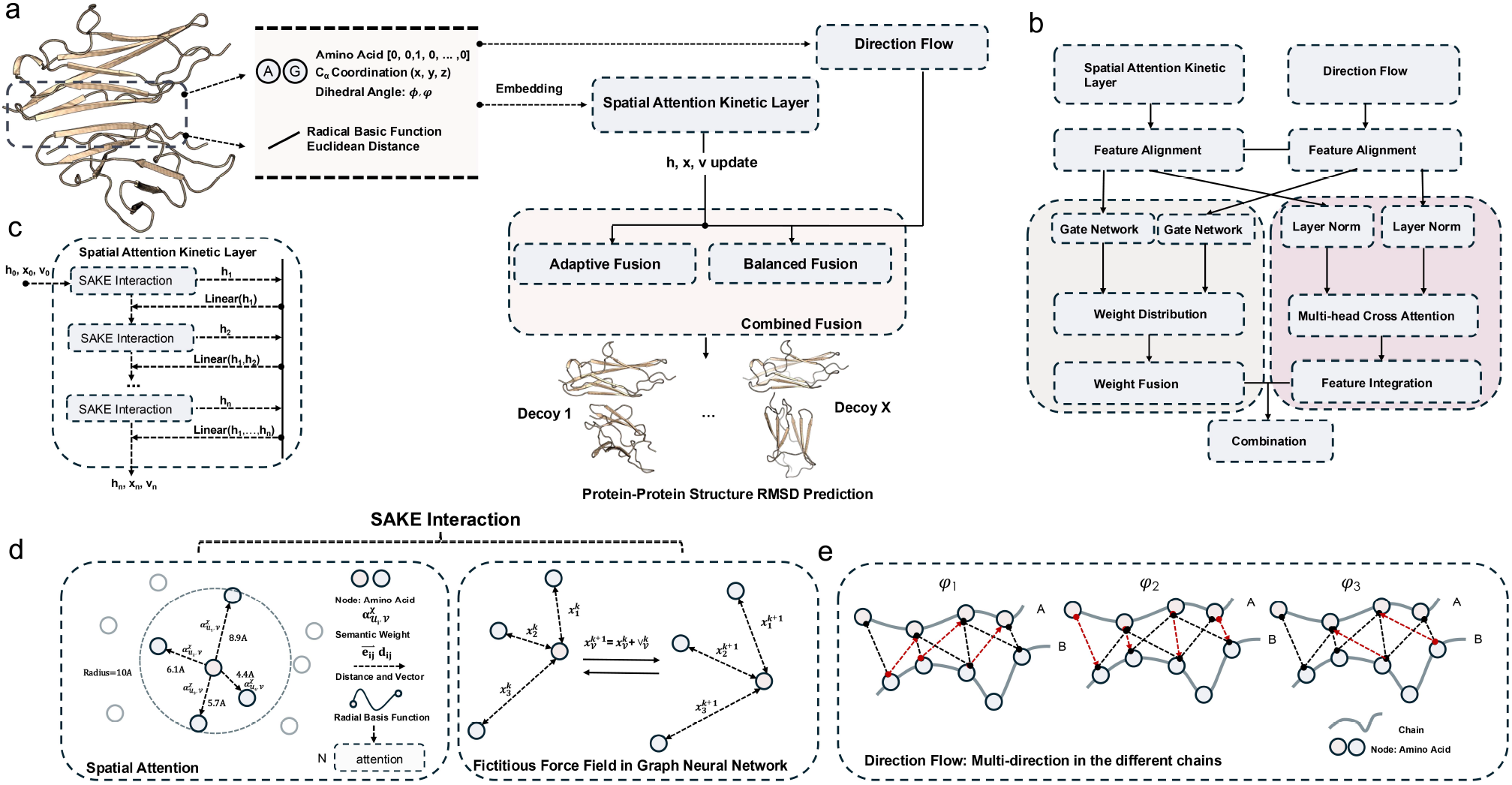
Workflow and architecture of SAKE-PP for protein–protein structure iRMSD prediction. (a) Workflow of SAKE-PP: Each protein complex is encoded using amino acid type, dihedral angles (*ϕ, ψ*), C*α* coordinates (*x, y, z*), and Euclidean distance–based features within a 10 Å interface region. The representations are passed to both the Direction Flow and the Spatial Attention Kinetic Layer. Their outputs are integrated in the Combined Fusion module to predict a scalar iRMSD value. (b) Feature fusion mechanism: The two representation streams undergo feature alignment and are processed by gate networks, weight fusion, and multi-head cross attention to yield a unified latent representation. (c) Spatial Attention Kinetic Layer: Composed of stacked SAKE Interactions, each connected via residual links. Each SAKE layer updates node features **h**, positions **x**, and velocities **v** in an *E*(*n*)-equivariant manner. (d) SAKE Interaction Module: Integrates spatial attention and a fictitious force field for joint node and coordinate updates. The spatial attention mechanism combines radial basis encoding of pairwise distances with edge-based semantic attention, capturing local geometric dependencies. A mixed attention score 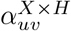 modulates neighborhood aggregation. Position updates are coupled with learned velocity vectors that evolve under a physics-inspired force field: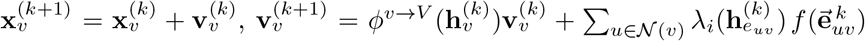. (e) Direction Flow: Captures multi-directional spatial interactions across protein chains using chain-aware angular relationships (*φ*_1_, *φ*_2_, *φ*_3_) to guide geometric feature propagation.

## Discussion

In this study, we introduce SAKE-PP, a GNN model specifically designed for evaluating protein–protein interfaces. The model consistently outperforms the native ranking score function of AF3 across multiple evaluation metrics, particularly excelling in its ability to discriminate between structural conformations. Our findings reveal a critical bottleneck in current structure prediction workflows: although deep learning models such as AF3 are capable of generating diverse and high-quality decoy ensembles, their internal confidence scores often fail to identify conformations with the most biologically meaningful minimum-energy states.

We further observe that AF3 frequently favors structurally deviated conformations over near-native alternatives—even when such alternatives exist within the generated ensemble—a tendency largely driven by inflated confidence scores. This systematic bias not only hampers accurate assessment of interfacial energetics but also undermines structure-based drug design efforts that rely on high-affinity binding predictions.

SAKE-PP addresses these limitations through a physics-inspired architecture that integrates spatial attention mechanisms with Laplacian eigenvector orientation, enabling more accurate representation of interfacial features. In cross-dataset evaluations, SAKE-PP achieves a 13.75% improvement in iRMSD-based conformation selection accuracy, indicating its superior capability in recovering experimentally determined structures relative to AF3 ranking score.

During training, we encountered several challenges. Although hierarchical sampling strategies were employed to capture diverse structural distributions, the model occasionally exhibits deviations between predicted and actual iRMSD values when encountering inputs far outside the training domain. This highlights the inherent complexity of modeling protein–protein interactions. Nevertheless, SAKE-PP maintains robust discriminative ability in fine-grained structural details, as particularly evident in its zero-shot generalization to antibody–antigen complexes: in this task, SAKE-PP outperformed AF3 ranking score by a correlation margin of 0.4, despite AF3 ranking score having been trained on a large quantity of antibody–antigen data. This result suggests that SAKE-PP captures fundamental physical principles governing protein interfaces rather than simply learning statistical patterns.

However, we also found that models relying solely on geometric features may sometimes diverge from biologically preferred conformations. For instance, in crystal structures derived from specific biological contexts such as signaling pathways, SAKE-PP selected a more tightly bound conformation, while AF3 ranking score favored a near-native pose. This discrepancy may arise from the absence of contextual biological information during training. Future models should consider incorporating biological priors or functional annotations to improve context-aware prediction.

To further validate these observations, we performed MD simulations on the protein complex 8PE2, comparing competing conformations selected by different scoring functions. Surprisingly, despite significant structural differences at the outset, all conformations remained stable throughout the simulation. This suggests the existence of multiple coexisting, energetically viable states in the protein–protein interaction landscape, offering a plausible explanation for why structurally distinct yet biophysically reasonable conformations may be selected by different models. Importantly, our binding energy analyses consistently demonstrated that SAKE-PP-selected conformations exhibit superior thermodynamic stability compared to AF3 ranking score-selected alternatives, even in cases where AF3 ranking score’s selection was closer to the reference crystal structure. This reinforces the model’s ability to identify energetically favorable binding modes that serve as better starting points for downstream applications, especially in finding drugs with high affinity.

Taken together, these findings highlight the necessity of ensemble-based strategies that incorporate multiple starting conformations to fully characterize protein complex interfaces. Looking forward, the SAKE-PP framework can be extended to more complex biomolecular assemblies, including protein–nucleic acid complexes and higher-order oligomers. Moreover, its integration with free energy calculations and adaptive sampling techniques holds promise for enhancing both accuracy and efficiency in structure prediction and drug discovery workflows.

## Methods

### Data preparation and preprocessing

Protein–protein complexes containing two distinct polypeptide chains were identified in the PDBBind dataset. ^32,33^ Structures were then refined using pdb4fixer by removing water molecules and other hetero atoms (e.g., small molecules), retaining only the main protein atoms. In cases where one chain was substantially shorter than the other, the shorter chain was designated as the “ligand,” while the longer chain was labeled the “receptor.” This procedure yielded 736 protein–protein complex structures in total.

Each protein–protein complex was docked using ZDock, generating 2000 conformations per complex.^18^ To introduce prior knowledge and ensure a reasonable docking space, a distance restraint of 15 Å was imposed between the “ligand” and “receptor,” based on their original closest contact. This restraint mimics the practical insight often employed in biochemical experiments to confine the search region around the putative binding site. Under this setting, a total of 1,472,000 candidate docking conformations were produced across all complexes.

In the evaluation of protein-protein docking conformations, we employed a hierarchical sampling strategy centered on the iRMSD. This approach was guided by prior findings: DiffDock ^34^ reports an average conformational distribution iRMSD of 4.85 Å, while CAPPI considers 4 Å, as the threshold distinguishing near-native conformations. Based on these observations, we designated the 4–5 Å range as a critical reference for assessing conformation quality.

Although a regression-based method was utilized during training, an adaptive sampling scheme was devised to ensure a balanced set of training samples, with a particular focus on the 4–5 Å range. This scheme addresses the typically uneven and non-normal distribution of docking conformations generated by ZDock across different protein complexes. Specifically, we calculated multiple metrics, including iRMSD, the fraction of native contacts (f_nat), and DockQ, to comprehensively evaluate the quality of all candidate conformations. The hierarchical sampling strategy was implemented as follows:If sufficient high-quality conformations were available within the 0–5 Å range, incremental sampling was performed at 10% intervals, complemented by uniform sampling in the 5–8 Å range to ensure representativeness. When the 0–5 Å range contained an insufficient number of conformations, the 10 highest-quality conformations from the 0–8 Å range were selected first, followed by uniform sampling of the remaining conformations until a total of 20 conformations was reached. In extreme cases, where the total number of conformations within the 0–8 Å range was fewer than 20, all available high-quality conformations were selected. Including the native conformations, we obtained a total of 15,456 conformations as our complete dataset.

Fig b illustrates the conformational distribution obtained after the application of a strategic sampling approach. According to the CAPRI classification criteria (with a threshold of 4 Å), the proportion of acceptable conformations is 42.87%, while the proportion of unacceptable conformations is 57.13%. Besides, among the acceptable conformations, high-quality conformations (RMSD 0–2 Å) account for 34.01%, whereas borderline acceptable conformations (RMSD 2–4 Å) constitute 65.99%. Figure c illustrates the distribution of 1,472,000 candidate docking conformations within the range of 0–10 Å. The distribution exhibits a sandglass-like pattern, with a relatively higher number of conformations at high RMSD values. After sampling, the distribution is transformed into the pattern shown in Figure d. Our sampling strategy ensures that the dataset achieves broad coverage of conformational distributions while maintaining a high level of sample representativeness in the critical range (RMSD 0–4 Å).

### Protein Structure Representation

We represent a protein structure as a graph 𝒢 = (𝒱, ℰ), where each node *v* ∈ 𝒱 corresponds to one residue in the protein backbone, and edges *E* capture their spatial proximity and biochemical relationships. Let |𝒱| = *N* denote the number of residues (nodes), and let ℰ ⊆ {(*u, v*) | *u, v* ∈ 𝒱} be the edge set.

#### Node Representation

For each residue *v* ∈𝒱, we collect both *spatial* and *chemical* features:

#### Spatial Features

- **C***α* **coordinates**: We treat the entire protein–ligand complex (or multi-chain complex) as a single structure and extract all C*α* atoms from the backbone. Each C*α* atom is indexed by a node *v*, whose 3D position is given by **x**_*v*_ ∈ ℝ^3^ (i.e., the (*x, y, z*) coordinates). These coordinates allow the model to capture the overall geometry of the protein complex.
- **Backbone dihedral angles**: For each residue *v*, we calculate the main-chain dihedral angles (*ϕ*_*v*_, *ψ*_*v*_) following the standard protein geometry definition. In rare cases where numerical issues arise (e.g., a tiny subset of angles failing to compute properly), we assign a small default value (e.g., 10^*−*5^) to avoid undefined behaviors in subsequent computations.
- **Angle-derived embedding**: We adopt the angle-derived embedding **a**_*v*_ ∈ℝ^3^ to convert dihedral angles into a 3D direction vector. Formally,

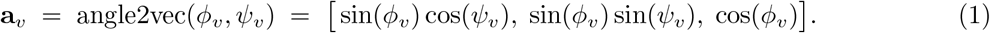

#### Chemical Features

- **Residue type encoding**: We represent each canonical amino acid with a one-hot vector or learned embedding, **r**_*v*_ ∈ℝ^20^.
- **Chain identifier**: Each residue *v* is associated with a chain identifier *c*_*v*_ ∈ℕ, which indicates which protein chain the residue belongs to (e.g., chain A, chain B, etc.).
- **C***α* **flag**: We include a binary flag *f*_*v*_ ∈ *{*0, 1*}* to specify whether the entry corresponds to a C*α* atom (main-chain *α* carbon).

By concatenating the chemical features above with the spatial features described previously, we arrive at the *complete node feature* vector. Specifically, we form

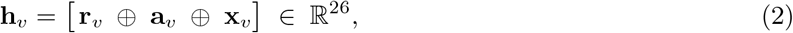

where *⊕* denotes vector concatenation. In this construction,

- **r**_*v*_ ∈ ℝ^20^ encodes the residue identity,
- **a**_*v*_ ∈ℝ^3^ (angle-derived embedding) represents dihedral angles,
- **x**_*v*_ ∈ℝ^3^ stores the C*α* coordinates.

Hence, each residue *v* is ultimately represented by a 26-dimensional feature vector that integrates both chemical and spatial information. This design aligns with our code’s practice of combining residue type, angle-based features, and 3D coordinates into a single node embedding.

#### Edge Representation

To fully capture inter-residue interactions, we define both the *edge set* ℰ and the *edge features* 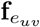 in one integrated procedure. Specifically, we first construct edges based on complementary criteria, then compute a rich set of geometric and biochemical descriptors for each edge.

#### Edge Construction

We adopt two strategies to determine whether an edge should exist between residues *u* and *v*:

1. **Contact-based rule**:

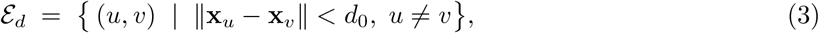

where *d*_0_ = 10 Å is a typical cutoff for identifying residue-residue contacts.
2. **k-Nearest Neighbors (kNN)**: For each residue *v*, we pick *k* = 30 closest neighbors by Euclidean distance on C*α* coordinates:

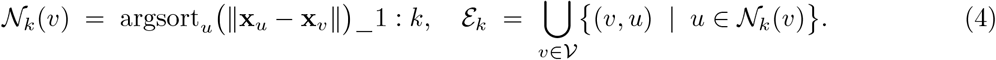

We then merge these sets to form the final edge set:

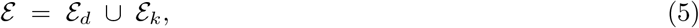

ensuring that both long-range contacts (within *d*_0_) and local neighbors (*k* = 30) are included.

#### Edge Features

Once the edge set ℰ is established, we compute an *edge feature vector* 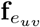 for each (*u, v*) ∈ ℰ. This vector encodes geometric and biochemical information essential for downstream message passing. Concretely, we combine:

- **Displacement vector and Distance**

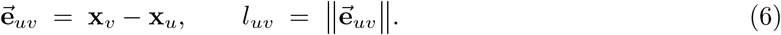
- **Direction**

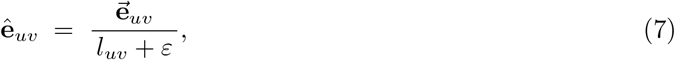

where *ε* is a small positive constant (e.g., 10^*−*6^) to avoid division by zero.
- **Radial Basis Expansion**: *I* n our implementation, we expand the raw distance *l*_*uv*_ via a set of radial basis functions:

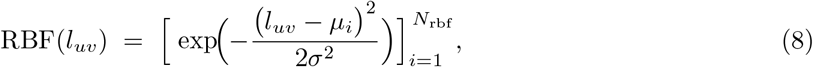

where 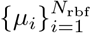 are centers evenly spaced in [0, *d*_0_], *σ* is a bandwidth, and we fix *N*_rbf_ = 16. This 16-channel representation provides a smooth and more expressive encoding of inter-residue distances.
- **Source and Target Residue Types**: For each edge (*u, v*), we can include the amino acid encodings **r**_*u*_, **r**_*v*_ ∈ ℝ^20^, thereby allowing the model to learn pairwise interaction patterns (e.g., hydrophobic vs. hydrophobic).
- **Chain Interaction Flag**

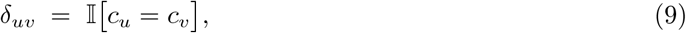

which indicates whether residues *u* and *v* belong to the same chain. In interface-focused tasks, one might specifically highlight the inter-chain edges (i.e., *c*_*u*_ *≠c*_*v*_).

Hence, the final edge feature vector (concatenating all the above) is:

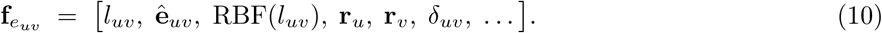

### SAKE Interaction Layer: Spatial Attention and Kinetic Updates

In the previous work, we present a noval neural network named **SAKE** (*spatial attention kinetic network with E(n)-Equivariance*) layer aims to simultaneously update the features of *(i)* nodes (residue embeddings) and *(ii)* their 3D positions with E (n) -equivariance. ^35^ In the following, we outline the essential components of SAKE and explain how they integrate into a single framework.

#### Spatial Attention Definition

Given a node *v* with embedding **h**_*v*_ ∈ ℝ^*C*^ and position **x**_*v*_ ∈ℝ^*n*^ in graph 𝒢, the *spatial attention ϕ*^SA^ is designed to capture geometric relationships among neighbors *u* ∈ 𝒩 (*v*):

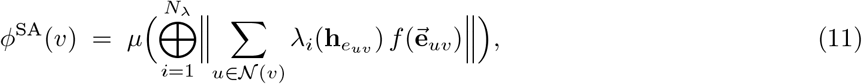

where 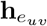 is the edge embedding, *λ*_*i*_ are attention weight functions (*i*.*e*., small MLPs) mapping edge embeddings to scalars, and *f* is an E(n)-equivariant transformation on the displacement vector 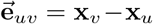.

#### Edge Embedding Update

Prior to applying attention weights, each edge (*u, v*) is assigned an embedding 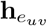 that combines both chemical and geometric features:

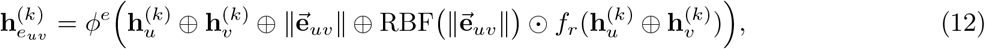

where RBF encodes distance into radial basis channels, and *f*_*r*_ is a small network that modulates the combined embeddings.

**Mixed Attention**. SAKE uses a combination of:

- A distance-based cutoff 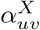 to smoothly suppress edges beyond a certain threshold *d*_0_
- A semantic attention 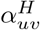 akin to GAT-style softmax weights that highlight edges with higher feature similarity

These are combined as:

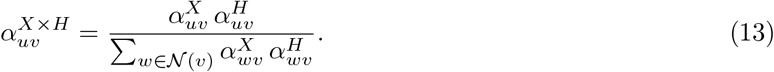

**Velocity and Position Updates**. The position updates are coupled with velocity vectors:

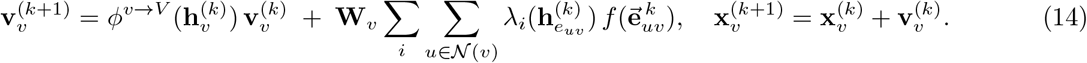

### Laplacian Eigenvector Directional Graph Neural Network

#### Laplacian Matrix and Eigenvectors

For an undirected graph 𝒢 = (*V, E*), we define the graph Laplacian matrix **L** ∈ ℝ^*N ×N*^ either in the combinatorial form **L** = **D** *−* **A** or in the normalized form 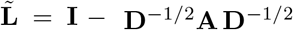, where **A** is the adjacency matrix (with *A*_*ij*_ = 1 if (*i, j*) ∈ *E* and 0 otherwise) and **D** is the degree matrix (*D*_*ii*_ = ∑*j A*_*ij*_). The Laplacian is symmetric positive semidefinite, so it admits an eigendecomposition

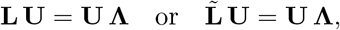

where **U** = [*ϕ*_0_ *ϕ*_1_ … *ϕ*_*N−*1_] is an orthonormal basis of eigenvectors, and **Λ** = diag(*λ*_0_, *λ*_1_, …, *λ*_*N−*1_) are the corresponding eigenvalues (sorted as 0 *≤ λ*_0_ *≤ λ*_1_ *≤* …). The first eigenvector *ϕ*_0_ associated with *λ*_0_ = 0 is the constant vector; The trivial eigenvector *ϕ*_0_ associated with the constant vector is discarded, and the subsequent k *nontrivial* eigenvectors *{ϕ*_1_, …, *ϕ*_*k*_*}* are retained to capture more meaningful structural properties of G.

#### Laplacian Eigenvector and directional aggregators in Laplacian Eigenvector Directional Graph Neural Network

Let 𝒢 = (*V, E*) be the input graph with |*V* | = *N* nodes and |*E*| = *M* edges. Denote the node features by 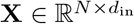 and the edge features by 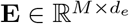. We first map **X** to the hidden dimension *d*_hid_ via

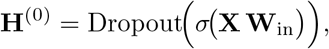

where 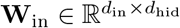 is a learnable matrix, and *σ* is the LeakyReLU activation function.

We compute the first *k* = 3 Laplacian eigenvectors **Φ** ∈ ℝ^*N ×*3^, serving as position embeddings (PE).

That is, each row **Φ**_*i*,:_ stores the eigenvector values (*ϕ*_1_(*i*), *ϕ*_2_(*i*), *ϕ*_3_(*i*)) for node *i*.

Specifically, for an edge (*i, j*), let *ϕ*_*k*_ denote the *k*th Laplacian eigenvector. The directional difference *F*_*i,j*_ = *ϕ*_*k*_(*j*)*−ϕ*_*k*_(*i*) encodes how node *j* differs from node *i* in the *k*th eigenvector dimension. Two specialized aggregators, *directional average* and *directional derivative*, utilize |*F*_*i,j*_| or *F*_*i,j*_ respectively to weigh neighbor features **h**_*j*_. In our work, these directional aggregators are combined with standard ones such as `mean`, `sum`, and `max`, along with degree-based *scalers* (`identity`, `amplification`, `attenuation`), to capture both global and local structures. Hence, DGNConv offers richer geometric expressiveness than purely isotropic graph convolutions.

#### Laplacian Eigenvector Directional Graph Neural Network Layers

For each layer *ℓ* = 1, …, *L*, we apply a Laplacian Eigenvector Directional Graph Neural Network module, denoted by *ℱ* ^(*ℓ*)^, which takes the graph (𝒢, **H**^(*ℓ−*1)^, **E, Φ**) as input. Inside *ℱ* ^(*ℓ*)^, we use five aggregators: `mean`, `max`, `sum`, `dir1-av`, `dir1-dx`, and three scalers: `identity`, `amplification`, `attenuation`. Combining each aggregator with each scaler yields multiple feature transforms that are concatenated and projected back to *d*_hid_. We then apply LeakyReLU and a dropout with rate *p* (e.g., *p* = 0.15). The output of layer *ℓ* is

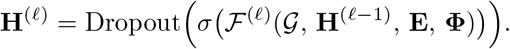

After stacking *L* layers of DGNConv, we obtain

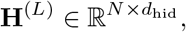

which we map to the output dimension *d*_out_ via a linear projection:

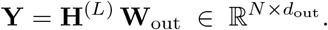

Here, 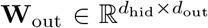 is a learnable parameter matrix. The returned node-level output **Y** thus has one row per node, each row being a *d*_out_-dimensional feature.

### SAKE-PP: Integrating SAKE and Laplacian Eigenvector Directional GNN

SAKE layers update both the node features and their 3D coordinates/velocities. We initialize the node features via a learnable linear map and optional activation/dropout,

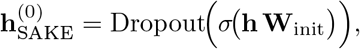

where 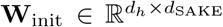. Then each SAKE Interaction Layer applies the spatial attention and kinetic updates (11–14), producing

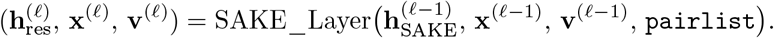

Each residual node embedding 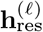 is then projected via an additional linear layer and concatenated with previous features, *e*.*g*.,

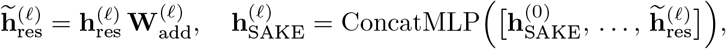

where ConcatMLP denotes a learned concatenation transform followed by batch normalization, activation, and dropout. After the final SAKE Interaction Layer (say *ℓ* = *L*_SAKE_), we obtain updated node embeddings 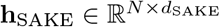, as well as new positions/velocities (**x, v**).

In parallel, original node features **h** and the edge features **E** (together with the graph structure 𝒢) are feeded into the Laplacian Eigenvector Directional GNN (Sec.). This module, denoted as `dgn`, outputs another node-level representation:

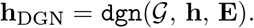

Internally, `dgn` uses multiple DGNConv layers, each exploiting both standard and directional aggregators (e.g. `dir1-av`, `dir1-dx`) with degree-based scalers, ultimately producing 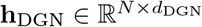.

We then concatenate the SAKE output **h**_SAKE_ and the DGN output **h**_DGN_ along the feature dimension:

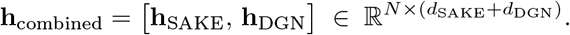

An additional learnable transformation `combined_processing` is applied, which may be a linear layer or a small MLP with activation:

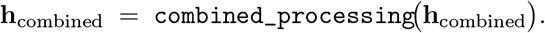

To obtain a graph-level representation, we perform global pooling over the node dimension:

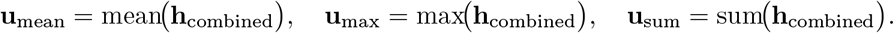

Here, mean, max, sum are taken node-wise (and separately within each subgraph if using a batched graph). We concatenate the three pooled vectors,

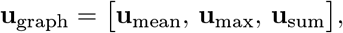

and finally project to the desired scalar output (or any target dimension) via

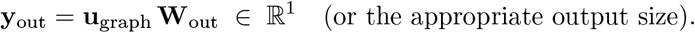

Thus, the `SAKE-PP` model jointly leverages the SAKE interaction mechanism (Eq. 11–14) and Laplacian Eigenvector Directional Aggregators to capture both local geometric relations (*via* positions and velocities) and global structural cues from the eigenvector-based directional GNN.

## Supporting information

SI for the manuscript including table and figure

## Data and Code Availability

All relevant data are provided in the Supplementary Information. The primary code could be found in the https://github.com/yxnyu/SakePP/.

## Conflicts of interest

There are no conflicts to declare.

## Acknowledgement

This study received partial support from the National Natural Science Foundation of China (NSFC); Grant Nos. 22333006, 92270001, 62376254, 32341017 and 32341018. Yuzhi Xu was funded exclusively by New York University. We gratefully acknowledge the high-performance-computing resources provided by NYU Abu Dhabi and the Greene supercomputing cluster at New York University, and we thank their dedicated staff for expert technical assistance.

## Footnotes

1 AF3 ranking score = 0.8 *×* ipTM + 0.2 *×* pLDDT

## Notes

### Competing Interest Statement

The authors have declared no competing interest.

