## Supplementary material for "Quantifying Protein-Protein Interaction with a Spatial Attention Kinetic Graph Neural Network": SI for the manuscript including table and figure

### 1 2024PDB Test-Set PDB IDs

8PE1, 8SW5, 8S61, 8XYZ, 8K0E, 9G5K, 9AZ4, 8YF2, 8YFT, 8RT9, 8W6B, 8PE2, 8S5W, 8RQC, 8QU7, 8K3A, 9FZD, 8EPH, 8SVE, 8S5U, 8RY2, 9B75, 8K4Q, 8IDA, 8R38, 8UW8, 8VEG, 8VTP, 9BHW, 8S1X, 8S4U, 9AXN, 8S4H, 8SJB, 8VGE, 8SFX, 8RTX, 8YFE, 8G4K, 8PMW, 8QVX, 8BOP, 8X7N, 9AXQ, 8WT4, 8RAJ, 9FZC, 8PUT, 8WZO, 8W6A, 8VVB, 8P6H, 8RPP, 8XF2, 8VK8, 8YSH, 8JZT, 8UMC, 8K9Z, 8FJ4, 8WBZ, 8XUP, 9GRQ, 9F98, 8X3G, 8VLT, 8QC1, 8QZ4, 9FDJ, 8GJR, 9G2A, 8SUY, 9FCJ, 9B76, 9FZK, 8Z5D, 8FRI, 8WEO, 8RO6, 9IY0, 8VK9, 9GNY, 9AXO, 8V00, 8Z8V, 8F2Z, 8RO9, 9D65, 9D6J, 8RO8, 8VGF, 9IUZ, 8EPC, 9D6K, 9D6M, 8Z5F, 9ATO, 9FW2, 8TYX, 8QXC, 9FZ4, 8YJ5, 8POX, 8RO7, 9BD3, 9BIY, 8F6S, 8QZ3, 8IK6, 8UG2, 8Q6Q, 9ASS, 8S0N, 8WIX, 8VJM, 8P2T, 8WPL, 9AVT, 8WK1, 8SOW, 9FEF, 8GDT, 8YOD, 8J09, 8RD3, 8VU4, 8VU1, 8VUC, 8WHZ, 8SFV, 8WBY, 8W70, 8KB8, 9GP2, 8ROH, 8WZN, 8SFS, 9B44, 9DP6, 8X77, 8PDC, 8WKS, 9B74, 8OWS, 8PEQ, 8WFH, 9F18, 8VTR, 8SLC, 8U70, 8TE7, 8IGD, 8ZJF, 8SG3, 8Z8M, 9ATP, 8PMZ, 8UX9, 8IKV, 8QJ0, 8PU3, 8T19, 8WFM, 8RZ0, 9IU1, 8VGG, 8W90, 8K33, 8JI9, 8XN2, 8XN5, 8Y0Y, 8UKV, 8SOZ, 9FO7, 8S4K

Most of these proteins were released after June 2024, ensuring they fall strictly outside the AlphaFold3 training window. Although we did not perform explicit homology filtering, this omission can only confer a limited advantage to AF3 ranking score, because the SAKE-PP training data are already subsumed within AF3 ranking score.

### 2 Antibody–Antigen Dataset PDB IDs

7QUH, 7ST5, 7TRH, 7U9E, 7UA2, 7UJA, 7UMN, 7UOW, 7UXL, 7WN2, 7WNB, 7XCZ, 7XDA, 7XDK, 7XDL, 7XEG, 7XEI, 7XIK, 7XIL, 7XJ6, 7XJ8, 7XJ9, 7XJF, 7XRZ, 7XS8, 7XSA, 7XSC, 7Y8J, 7YD1, 7YDS, 7YK4, 7YRU, 7YUE, 7YV1, 7ZOZ, 7ZQT, 8A44, 8A96, 8A99, 8AHN, 8AV9, 8BCZ, 8BSE, 8BSF, 8BYU, 8C3V, 8CIM, 8CT6, 8CWI, 8CWJ, 8CWK, 8CZ8, 8CZZ, 8D7E, 8D9Y, 8D9Z, 8DA0, 8DA1, 8DB4, 8DE4, 8DN6, 8DN7, 8DNN, 8DTO, 8DUZ, 8DWW, 8DWY, 8DY1, 8DY5, 8DYX, 8DZ3, 8DZV, 8E1G, 8E1M, 8E6J, 8E6K, 8EE0, 8EE1, 8EK1, 8EKA, 8EL2, 8ELO, 8ELP, 8ELQ, 8EOO, 8EPA, 8F0H, 8F60, 8F6L, 8F6O, 8FAX, 8FG0, 8G30, 8G3M, 8G3N, 8G3O, 8G3P, 8G3Q, 8G3R, 8G3V, 8G3Z, 8GB6, 8GB7, 8GB8, 8GNK, 8GS9, 8GTP, 8GTQ, 8H07, 8HC4, 8HC5, 8HEB, 8HEC, 8HED, 8HHX, 8HHY, 8HN6, 8HN7, 8I5H, 8I5I, 8IB1, 8IUK, 8IUM, 8IW9, 8OL9, 8SAQ, 8SAR, 8SAS, 8SAV, 8SAW, 8SAX, 8SAY, 8SB0, 8SB1, 8SB2, 8SB3, 8SB5, 8SCX, 8SMT

### 3 Molecular Dynamics Simulations

The complexes of 8K0E, 8YF2, 8S4K, 8SOZ, 8VGG, and MOD were predicted by both AF3 ranking score and SAKE-PP. All complexes were prepared using the leap module in AMBER20 package. All MD simulations were performed using pmemd.cuda in AMBER20 with ff14SB force field for the protein. All simulations were carried out in a truncated octahedron box formed by TIP3P explicit water molecules. The closest distance between any atom originally present in solute and the edge of the periodic box is 12 Å. The particle mesh Ewald (PME) method is used to treat the long-range electrostatic interactions. The nonbonded interactions are truncated with 10 Åcutoff. Periodic boundary condition (PBC) is imposed on the system during the calculation of nonbonded interactions. The time-step is set at 2 fs and SHAKE is used to constrain the bonds involving hydrogen atoms. Langevin thermostat with the collision frequency 2.0 is applied to control the temperature.

First, the system is minimized with protein constrained to equilibrate the solvent. Then, the protein is released to minimize the whole simulation system. The system is slowly heated to 300 K, followed by a 40 ns equilibration of the whole system in an NPT ensemble at an interval of every 10 fs, and during this section, no restraint is performed. Then a 10 ns simulation was performed to obtain 5000 snapshots for ASGBIE analysis.

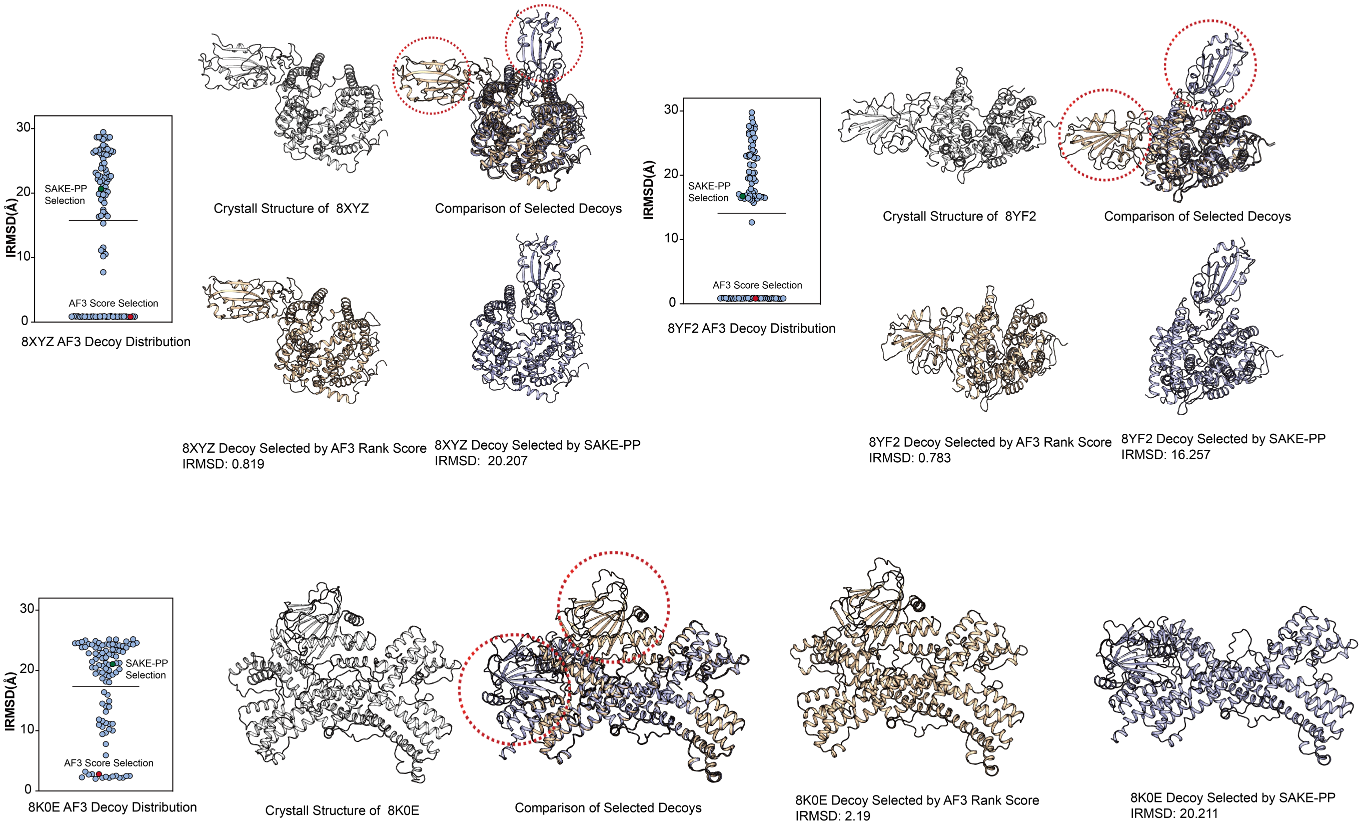


Figure S1: (a, d, k) Scatterplots of iRMSD for all 100 AlphaFold3 (AF3)–generated decoys of PDB IDs 8XYZ, 8YF2, and 8K0E, respectively. The green marker indicates the decoy chosen by SAKE-PP, while the red marker denotes the one selected by the AF3 ranking score. The grey horizontal line marks the iRMSD of the SAKE-PP selection. Percentages report the fraction of decoys whose iRMSD is equal to or lower than that of the AF3-selected model. (b, e, l) Crystal structures of the three targets, rendered in grey. (c, f, m) Structural overlays of the two selected decoys: the AF3 choice is shown in wheat, the SAKE-PP choice in blue. Red dashed circles highlight interface regions where the two selections diverge markedly. (g, i, n) Isolated views of the decoys selected by AF3, with their corresponding iRMSD values. (h, j, o) Isolated views of the decoys selected by SAKE-PP, also annotated with their iRMSD values.

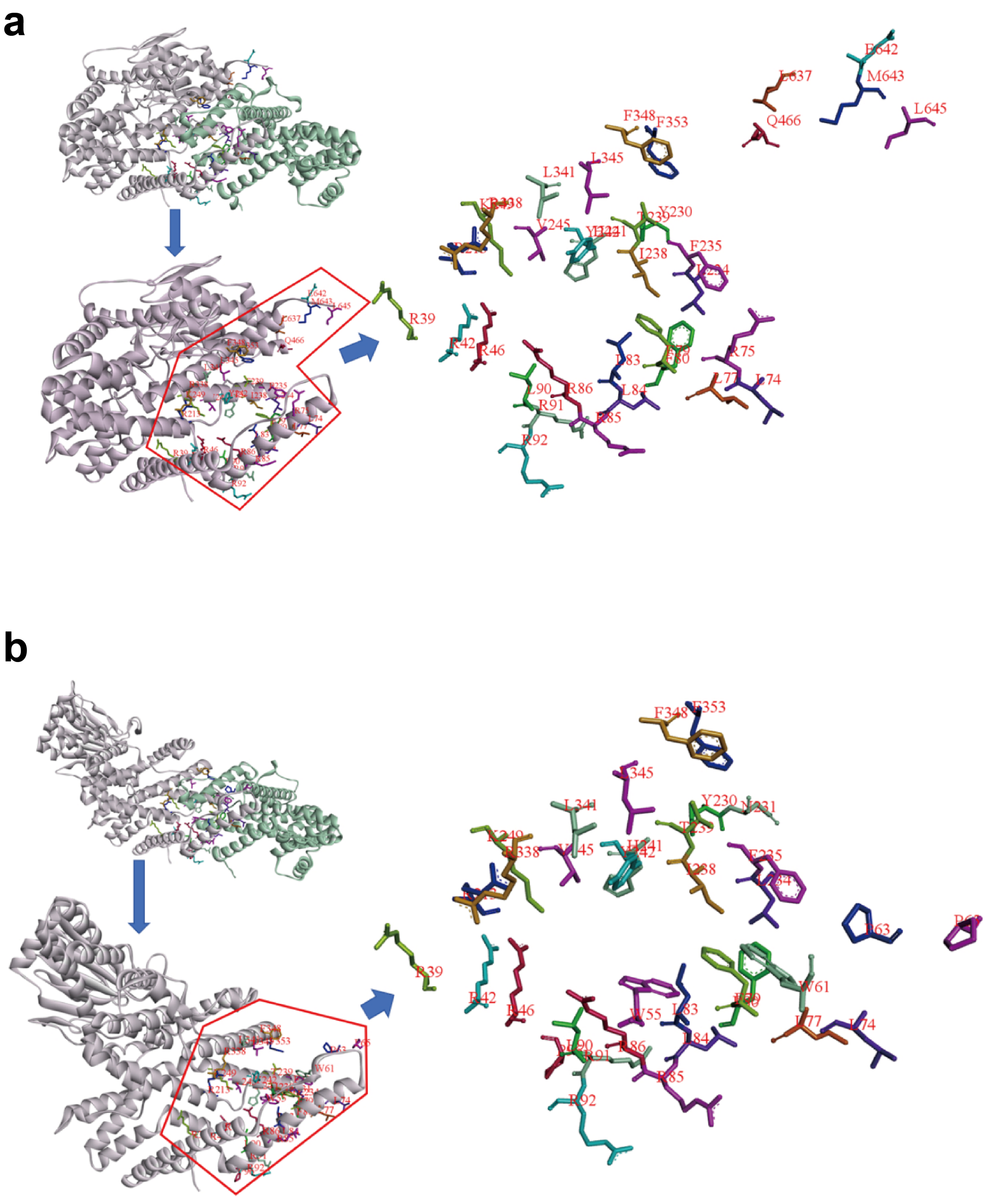


Figure S2: Comparison of predicted interface hotspots in the 8K0E complex using two scoring methods: (a) hotspots identified by the AF3 ranking score; (b) hotspots identified by the SAKE-PP score.

Table S1: Per-residue ddG contributions for 8K0E using AF3 ranking score vs. SAKE-PP.

| **8K0E – AF3 ranking score** | | **8K0E – SAKE-PP** | |
|---|---|---|---|
| **Residue** | **ddG** | **Residue** | **ddG** |
| 242TYR | -6.71 | 91ARG | -6.53 |
| 91ARG | -6.41 | 242TYR | -6.05 |
| 86ARG | -6.22 | 92ARG | -5.41 |
| 77LEU | -5.31 | 77LEU | -5.05 |
| 84LEU | -4.83 | 84LEU | -4.95 |
| 234LEU | -4.12 | 235PHE | -4.73 |
| 353PHE | -4.10 | 86ARG | -4.01 |
| 235PHE | -3.68 | 83LEU | -4.00 |
| 238ILE | -3.52 | 353PHE | -3.79 |
| 241HIE | -3.38 | 234LEU | -3.78 |
| 74LEU | -3.11 | 241HIE | -3.67 |
| 345LEU | -3.03 | 345LEU | -3.42 |
| 341LEU | -3.01 | 238ILE | -3.10 |
| 338ARG | -2.84 | 74LEU | -2.95 |
| 85ARG | -2.75 | 341LEU | -2.80 |
| 466GLN | -2.73 | 80PHE | -2.50 |
| 80PHE | -2.55 | 338ARG | -2.47 |
| 83LEU | -2.23 | 46ARG | -2.31 |
| 348PHE | -2.04 | 85ARG | -2.16 |
| 90LEU | -2.03 | 79PHE | -2.14 |
| 230TYR | -1.99 | 61TRP | -2.02 |
| 79PHE | -1.80 | 65PRO | -1.96 |
| 643MET | -1.67 | 55TRP | -1.93 |
| 645LEU | -1.62 | 231ASN | -1.78 |
| 42ARG | -1.46 | 96PRO | -1.77 |
| 249LYS | -1.43 | 230TYR | -1.72 |
| 245VAL | -1.41 | 42ARG | -1.63 |
| 637LEU | -1.35 | 245VAL | -1.47 |
| 92ARG | -1.31 | 249LYS | -1.44 |
| 75ARG | -1.30 | 39ARG | -1.35 |
| 46ARG | -1.26 | 213ARG | -1.24 |
| 213ARG | -1.25 | 348PHE | -1.18 |
| 642GLU | -1.25 | 90LEU | -1.16 |
| 39ARG | -1.20 | 239THR | -1.10 |
| 239THR | -1.12 | 63PRO | -1.02 |

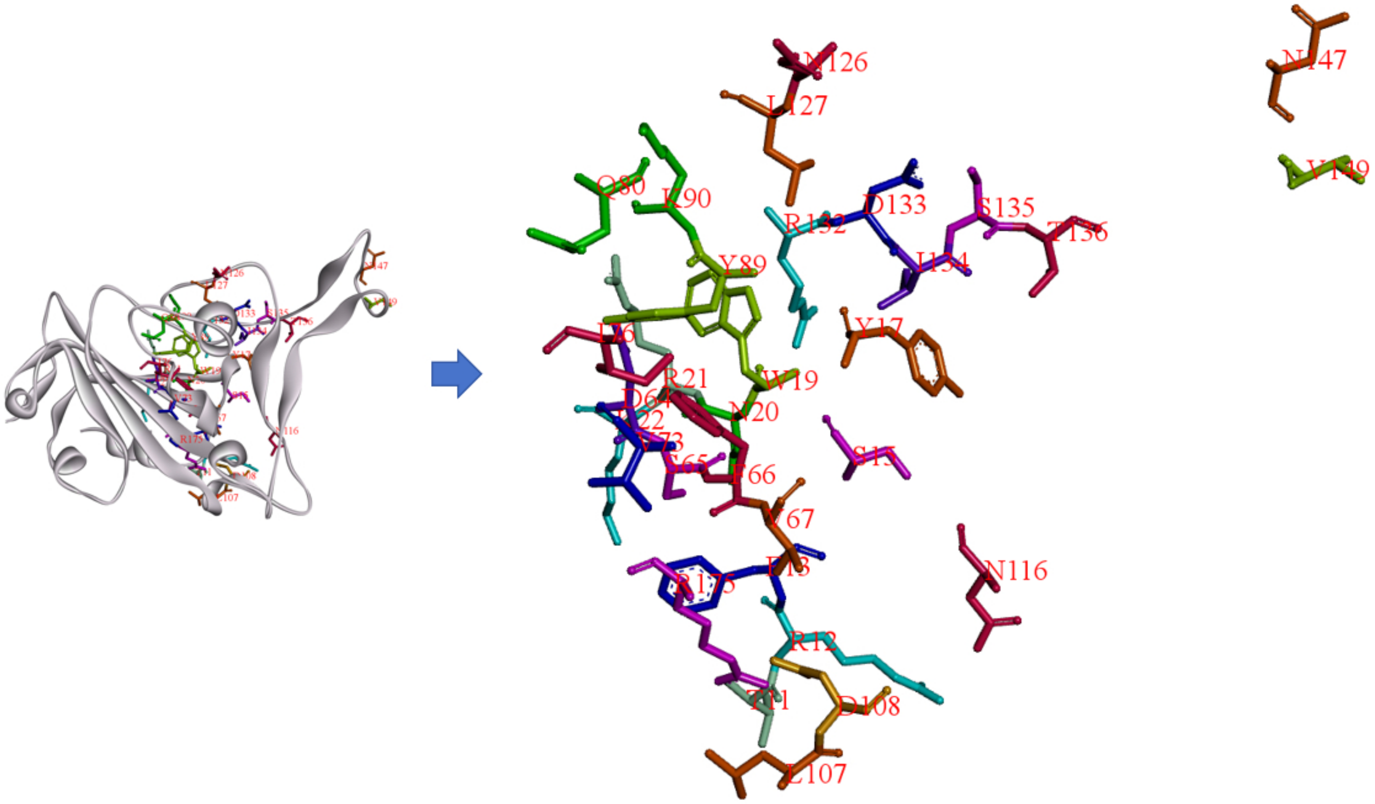


Figure S3: Comparison of predicted interface hotspots in the 8YF2 complex using two scoring methods.

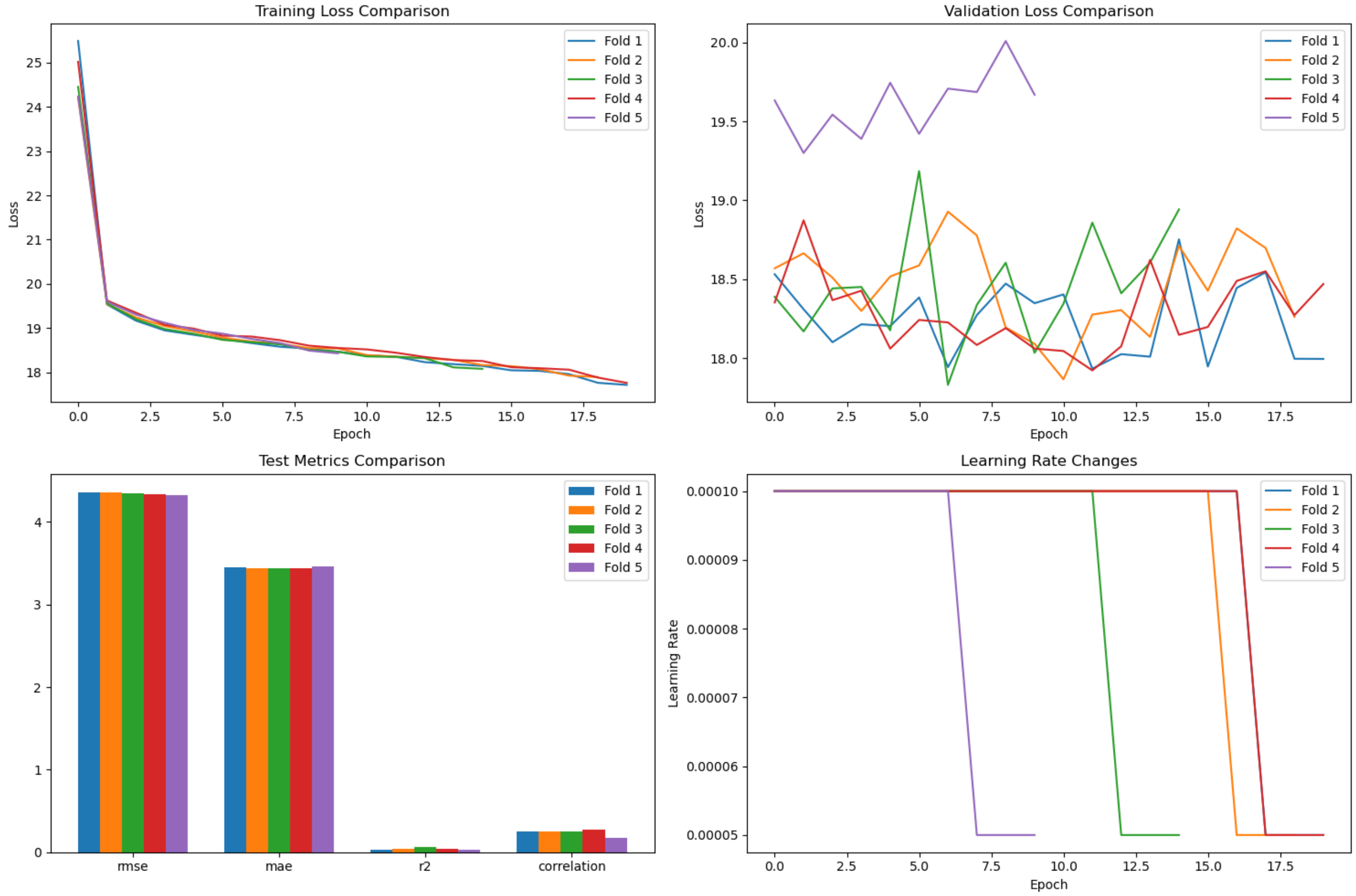


Figure S4: Cross-validation training dynamics and performance evaluation in None-strategy training. (a) Epoch-wise training loss MSE curves for each of the five folds; colors correspond to individual folds as indicated in the inset legend. (b) Validation-set loss trajectories for the same folds, highlighting inter-fold variability across epochs. (c) Test-set performance of each fold, reported as RMSE, MAE, coefficient of determination ($R^2$), and Pearson correlation; bars are color-coded by fold. (d) Learning-rate schedules used during training: all folds start at $1.0 \times 10^{-4}$ and undergo step-decays at fold-specific epochs (purple at epoch 6, green at 12, orange at 15, red at 17, and blue unchanged through 20 epochs).

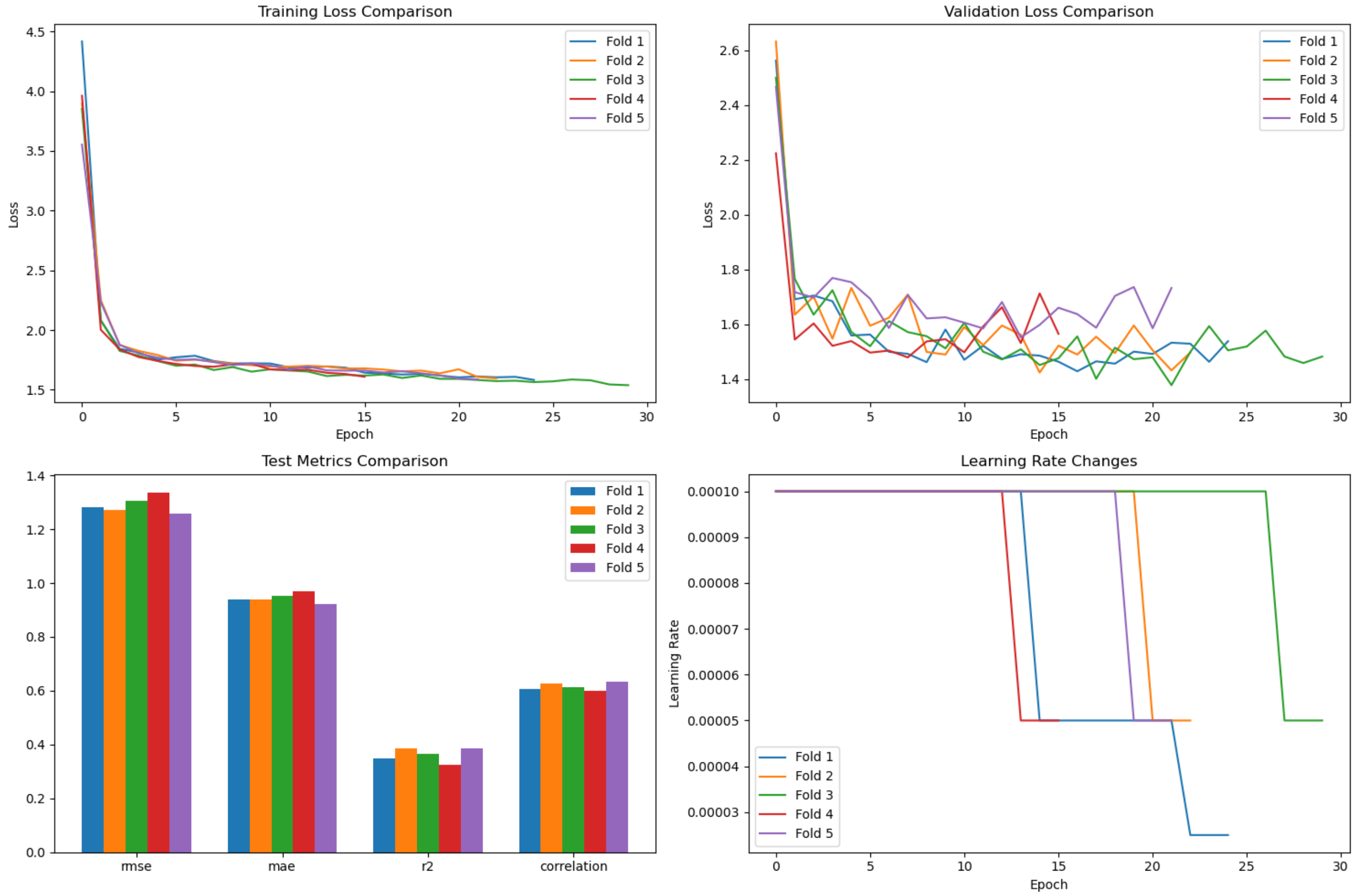


Figure S5: Cross-validation training dynamics and performance in the Strategy training. (a) Epoch-wise training loss (MSE) curves for the five folds; colours match the inset legend. All folds converge rapidly during the first few epochs and gradually plateau near 1.6. (b) Validation-set loss trajectories for the same folds, showing fold-to-fold variability yet an overall downward trend that stabilises around 1.6 by epoch 30. (c) Test-set evaluation of each fold, reported as RMSE, MAE, coefficient of determination ($R^2$), and Pearson correlation; bars are colour-coded by fold. (d) Learning-rate schedules: all folds start at $1.0 \times 10^{-4}$ and undergo two step-decays to $5.0 \times 10^{-5}$ and $2.5 \times 10^{-5}$ at fold-specific epochs (purple and red at ∼13–15, orange at 20, blue at 19/22, green at 26), illustrating adaptive optimisation across folds.
